# Structure of a bacterial ribonucleoprotein complex central to the control of cell envelope biogenesis

**DOI:** 10.1101/2022.01.04.474903

**Authors:** Md. Saiful Islam, Steven W. Hardwick, Laura Quell, Dimitri Y. Chirgadze, Boris Görke, Ben F. Luisi

## Abstract

The biogenesis of the essential precursor of the bacterial cell envelope, glucosamine-6-phosphate (GlcN6P), is controlled through intricate post-transcription networks mediated by GlmZ, a small regulatory RNA (sRNA). GlmZ stimulates translation of the mRNA encoding GlcN6P synthetase in *Escherichia coli*, but when bound by the protein RapZ, it becomes inactivated through cleavage by the endoribonuclease RNase E. Here we report the cryoEM structure of the RapZ:GlmZ complex, revealing a complementary match of the protein tetrameric quaternary structure to an imperfect structural repeat in the RNA. The RNA is contacted mostly through a highly conserved domain of RapZ that shares deep evolutionary relationship with phosphofructokinase and suggests links between metabolism and riboregulation. We also present the structure of a pre-cleavage encounter intermediate formed between the binary RapZ:GlmZ complex and RNase E that reveals how GlmZ is presented and recognised for cleavage. The structures suggest how other encounter complexes might guide recognition and action of endoribonucleases on target transcripts, and how structured substrates in polycistronic precursors are recognised for processing.

## Introduction

In all known bacteria, the post-transcriptional control of gene expression is fine-tuned and integrated into elaborate networks through the actions of numerous small regulatory RNAs (sRNAs). These RNA species can boost or suppress expression of target mRNAs to which they base-pair with imperfect complementarity (Wagner and Romby, 2015; Hör et al., 2020). sRNA activities are mediated by RNA binding proteins that faciliate specific and regulated recognition, and they can guide globally acting ribonucleases to silence specific targets. A few sRNA-binding proteins including the chaperones Hfq, ProQ and CsrA have been studied and structurally elucidated in bacteria (Holmqvist and Vogel, 2018; Babitzke et al., 2019; Quendera et al., 2020). However, how such RNA-chaperones act mechanistically to achieve selective recognition and turnover of the target RNA by the ribonuclease is still unsolved.

A salient example of sRNA-mediated riboregulation in network control is bacterial cell envelope biogenesis. A key component of the cell envelope is a peptidoglycan layer that provides mechanical robustness and cellular integrity. In addition, Gram-negative bacteria are surrounded by outer membranes containing lipopolysaccharide (LPS), which are recognized by the host innate immune system, but also provide protection against many antimicrobials including antibiotics. Biosynthesis of both the peptidoglycan layer and LPS relies on UDP-GlcNAc, which is produced from the amino-sugar glucosamine-6 phosphate (GlcN6P) (Khan *et al*., 2016). GlcN6P can be generated *de novo* by the enzyme GlmS but may also derive from recycling of the cell wall and available exogenous amino sugars. When cells grow rapidly or encounter antimicrobials that inhibit GlmS, the ensuing deficiency of GlcN6P is sensed to trigger a boost in synthesis of GlmS mediated by a post-transcriptional regulatory network, and this ensures that the supply of the required metabolite meets the cellular demand (Khan et al., 2016).

Cellular levels of GlcN6P are sensed by the sRNA-binding protein RapZ (Khan et al., 2020) (Figure 1). Low levels of the metabolite favour RapZ interaction with a two-component signalling system formed by the proteins QseE and QseF. This triggers expression of the sRNA GlmY, which in turn is sequestered by RapZ to form a stable complex. RapZ can also bind a structural orthologue of GlmY, known as GlmZ (Figure 1). High levels of GlmY displace GlmZ, which associates instead with the RNA chaperone protein Hfq. Hfq binding to GlmZ protects the sRNA and facilitates its base-pairing with an Anti-Shine-Dalgarno sequence in the *glmS* mRNA encoding glucosamine-6-phosphate synthase. This interaction boosts translation of the mRNA, increasing the levels of the enzyme and the metabolite (Kalamorz *et al*., 2007; Urban and Vogel, 2008; Göpel *et al*., 2013; Göpel *et al*., 2016; Reichenbach et al., 2008). When GlcN6P levels are sufficiently high, GlmY is released by RapZ and degraded, allowing RapZ to bind GlmZ instead (Khan et al., 2020). Ultimately, RapZ presents GlmZ to the endoribonuclease RNase E in a manner that directs cleavage within the base pairing site of the sRNA, thereby inactivating it and silencing synthesis of the synthase (Göpel *et al*., 2013; Durica-Mitic *et al*., 2020; Gonzalez et al., 2017; Figure 1).

**Figure 1.**
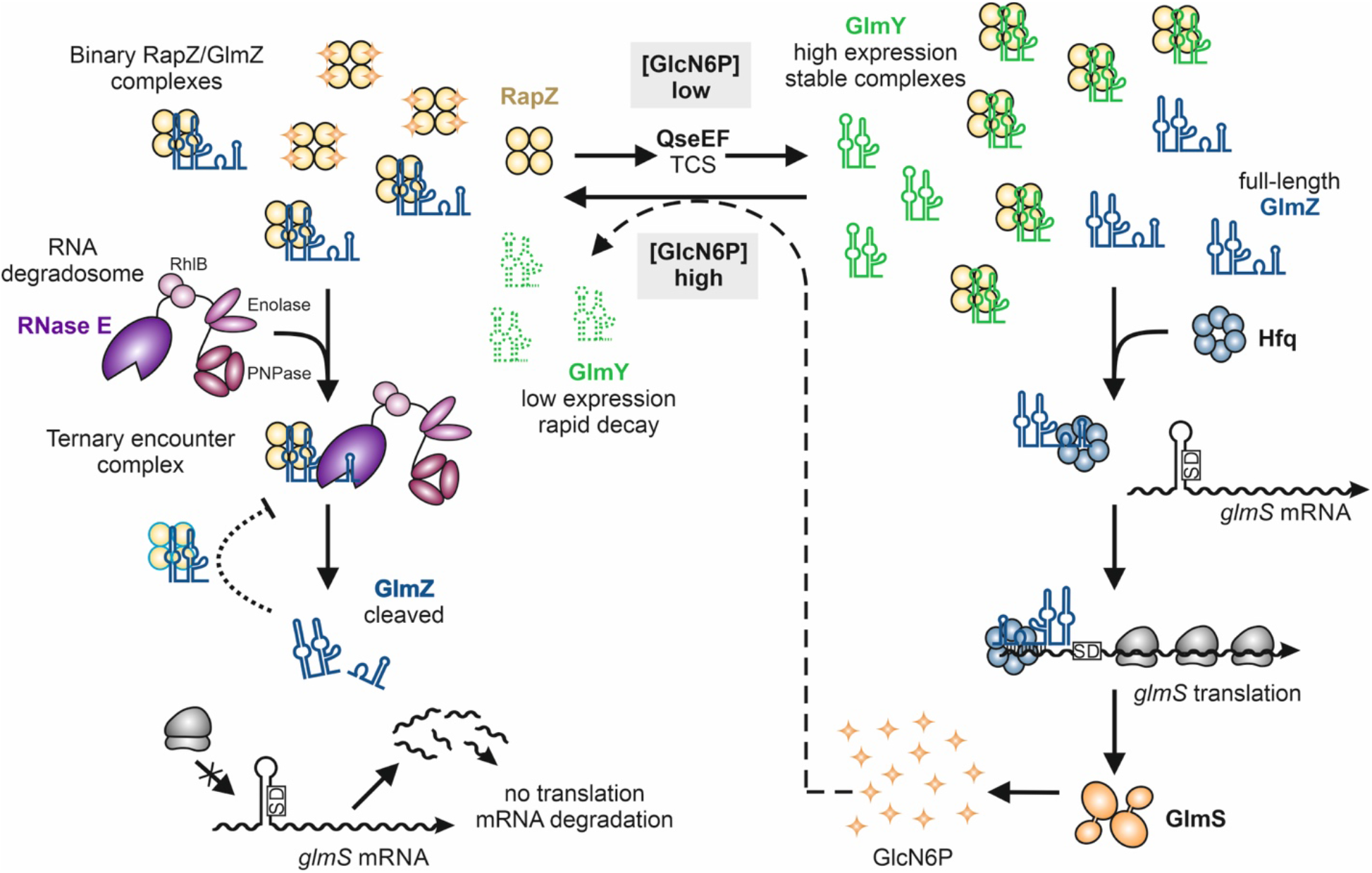
Model illustrating the role of RapZ for small RNA-mediated regulation of GlcN6P synthesis (according to Khan et al., 2020). When sensing a low level of GlcN6P, RapZ boosts expression of the sRNA GlmY via interaction with a two-component system. Elevated GlmY levels sequester RapZ into stable complexes. This allows the homologous sRNA GlmZ to interact with Hfq, which promotes base-pairing with the *glmS* mRNA. GlmZ base-pairing stimulates *glmS* translation, thereby replenishing GlcN6P through elevated GlmS enzyme levels (right). When the GlcN6P concentration is sufficiently high, RapZ is released from complexes with GlmY and the sRNA is rapidly degraded. As a result, RapZ is free to bind and present GlmZ to RNase E, which cleaves the sRNA in the base-pairing site. Cleaved GlmZ is unable to stimulate *glmS* translation, resulting in basal enzyme levels (left).

RNase E is a highly conserved, hydrolytic endoribonuclease encoded by bacteria of diverse families, and the enzyme is known to prefer to cleave single-stranded regions (SSRs) at the phosphate located two nucleotides upstream from a uracil (Chao *et al*., 2017). One mechanistic puzzle is why RNase E requires RapZ to cleave GlmZ. Clues as to how other RNAs might be recognised by RNase E have been provided through insight into crystal structures of the catalytic domain of the *E. coli* enzyme in apo and RNA bound forms, revealing a homotetramer organised as a dimer-of-dimers and details of the active site (Figure S1a) (Callaghan et al., 2005; Bandyra et al. 2018; Koslover et al., 2008). Experimental data for RapZ show that it forms a homotetramer and that the fold of the protomer comprises two domains of roughly equal size (Gonzalez *et al*., 2017). The N-terminal domain (NTD) bears Walker A and B motifs and resembles an adenylate kinase, while the C-terminal domain (CTD), which is sufficient for RNA binding, bears structural similarity with the sugar binding domain of phosphofructokinase. The quaternary structure of the RapZ tetramer is distinctive, with N and C-terminal domains forming separate symmetrical dimers lying at the apexes of the RapZ tetramer in a cyclic organisation (Figure 2b). Molecular genetics data indicate the requirement of the quaternary structure for stimulation of RNase E cleavage activity *in vitro* and overall regulatory activity *in vivo* (Gonzalez *et al*., 2017; Durica-Mitic *et al*., 2020). It is likely that the tetrameric organisation is required to present GlmZ for recognition and silencing by RNase E.

**Figure 2.**
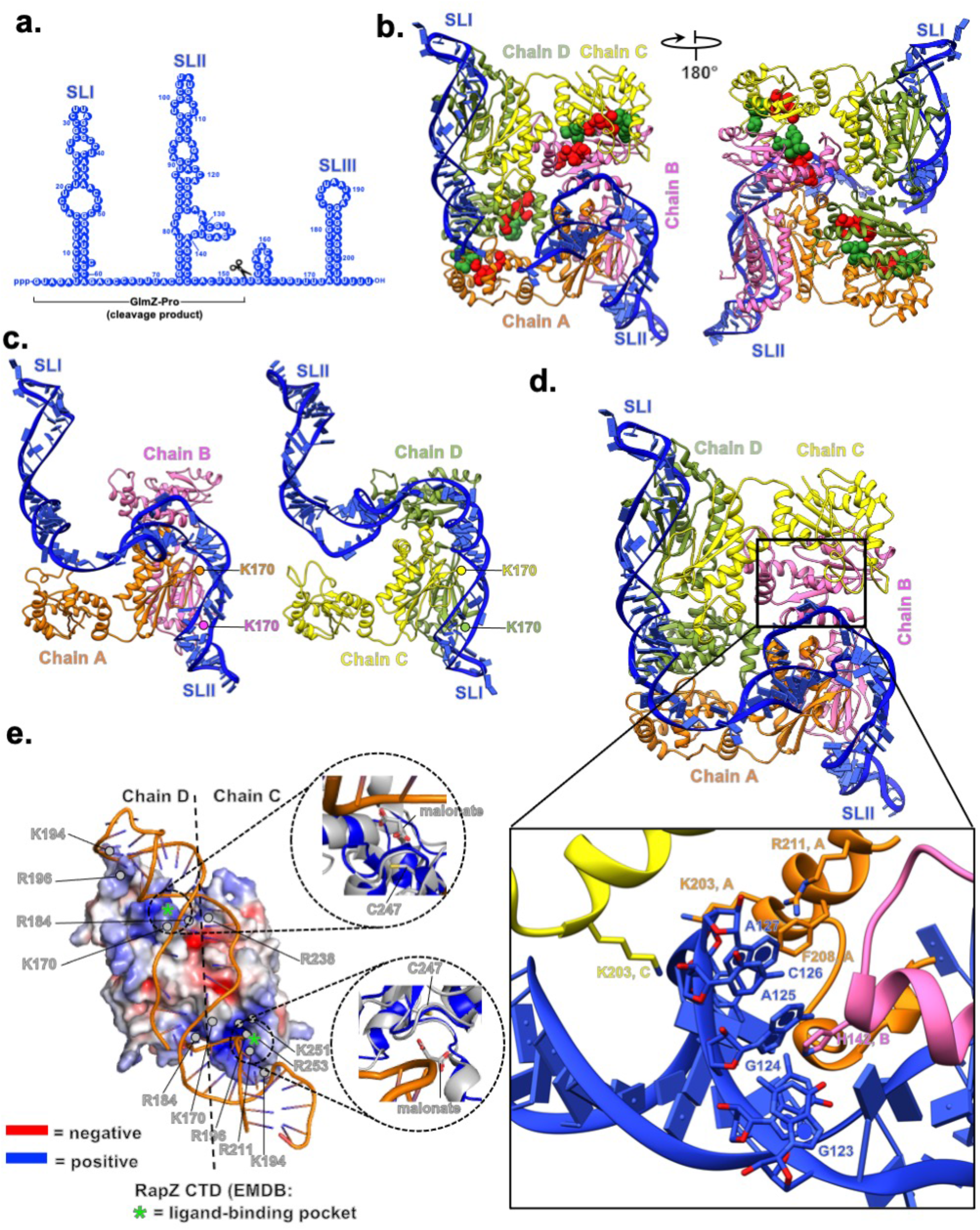
The cryoEM structure of the RapZ:GlmZ binary complex. (a) GlmZ secondary structure predicted by ViennaRNA Package 2.0 (Lorenz *et al*. 2011). (b) 3D model of the RapZ:GlmZ binary complex. The residues of the Walker A and B motifs are coloured red and green, respectively. (c) Similarity of the interactions of RapZ C-terminal domain (CTD) with SLI and SLII; a side-by-side comparison of the interactions of the stem loops with dimer sub-assemblies of RapZ. (d) Interaction between SLII and residues of RapZ chains A, B, and C. K203^A,C^ interacts with the phosphate backbone; F208^A^ interacts with A127^GlmZ^, and H142^B^ interacts with A125^GlmZ^. (e) View down the pseudo-fold axis of SLI interacting with a dimer of the RapZ CTD. The RNA duplex is underwound, which is apparent from the widened groove as viewed in this perspective. Interaction with GlmZ is achieved by the complementary electrostatic surface of the RapZ structure. The RapZ CTD is enriched mostly with basic residues. The insets show overlays revealing that the location of malonate molecules in the crystal structure of apo-RapZ (PDB:5O5S) occurs at the sites contacting GlmZ in the GlmZ:RapZ binary complex.

While detailed structures are available for apo-RapZ, there is no structural insight illuminating how the protein recognises GlmZ or GlmY. Secondary structure predictions and structural probing for GlmZ indicate 3 major stem loop (SL) structures (I, II, III) that in principle could provide key recognition features for both RapZ and RNase E (Göpel *et al*., 2013, 2016) (Figure 2a). SL III in GlmZ serves as transcriptional terminator, and does not appear critical for directing RNase E cleavage since it can be replaced with non-cognate terminators (Göpel et al., 2016). RapZ directs RNase E to cleave GlmZ at a site located in the SSR between SLII and SLIII (Figure 2a). Structural probing experiments did not indicate any structural re-organisation of GlmZ when bound to RapZ (Göpel *et al*., 2013), which suggests that the protein does not simply make the cleavage site accessible for RNase E through induced fit.

Here, we have used cryoEM to solve the structures of both the binary complex of RapZ:GlmZ and the ternary complex of the RNase E catalytic domain (RNase E-NTD), RapZ and GlmZ. The structures reveal how an imperfect structural repeat in the RNA matches the tetrameric quaternary structure of RapZ. The protein largely contacts the RNA through the phosphofructokinase-like domain, supporting the hypothesis that ancient metabolic enzymes can moonlight as RNA binders with regulatory consequences. Our data provide insights into the molecular recognition and the basis for discrimination between structurally similar RNAs through binding of imperfect RNA duplex repeats and presentation of a SSR to the catalytic site of RNase E. Our models suggest a general RNase E recognition pathway for complex substrates, and how other RNA chaperones such as Hfq might work in an analogous assembly to present base-paired sRNA/mRNA pairs for cleavage.

## RESULTS

### In vitro reconstitution of RapZ:GlmZ and RNase E:RapZ:GlmZ complexes

Previous reports have shown that, in the presence of RapZ, the catalytic domain of RNase E (RNase E-NTD) is sufficient to cleave GlmZ *in vivo* as well as *in vitro* (Göpel et al., 2013; Gonzalez et al, 2027; Durica-Mitic et al., 2020). To capture a pre-cleavage Michaelis-Menten state of the catalytic domain, aspartate 346 in the active site of RNase E-NTD was replaced with cysteine (Figure S1a), which changes the metal requirement for ribonuclease activity from Mg^2+^ to Mn^2+^ (Thompson, Zong and Mackie, 2015). An *in vitro* assay including RapZ confirms that this variant is inactive in the presence of Mg^2+^ but can cleave GlmZ in the presence of Mn^2+^ (Islam et al., 2021) (Figure S1b). We purified the recombinant mutant RNase E-NTD, RapZ and GlmZ and combined these components to reconstitute the ternary ribonucleoprotein complex (Figure S2). Reconstitution was performed in buffer without Mn^2+^ but with Mg^2+^to ensure proper folding of GlmZ and to lock the complex in the pre-cleavage encounter state. Formation of an equilibrium complex containing mutant RNase E-NTD, RapZ, and GlmZ was confirmed by size exclusion chromatography (SEC) and mass-photometry (Figure S2). Following extensive optimisation, cryo-EM datasets were collected for the RNase E-NTD:RapZ:GlmZ complex on grids coated with graphene oxide (Figure S3). Initial particle classification and map refinement revealed two main subclasses; one featuring the binary complex comprising one copy of GlmZ and one copy of RapZ tetramer (Figure S3), and a second class a ternary complex of RNase E-NTD:RapZ:GlmZ. The ternary complexes were heterogeneous and had either one or two RapZ:GlmZ associated with the RNase E-NTD (Figure S4). The ‘fundamental unit’, comprising one RNase E-NTD dimer, one RapZ tetramer and one GlmZ RNA (∼ 630 kDa) was refined through masking and particle substraction (Figure S3, right panel), as will be discussed further below.

### The RapZ:GlmZ binary complex and RNA structure recognition by the phosphofructokinase– like domain

A map of the binary complex was obtained at 4.4 Å resolution based on Fourier shell correlation (Figure S5, Table S1). This map provides an envelope that accommodates well a crystal structure of the RapZ tetramer (Gonzalez *et al*., 2017). Remaining density allowed for building of the first two stem loops of GlmZ RNA (SLI and SLII) (Figure 2a, 2b). The RNA spans over the surface of the RapZ tetramer forming interactions that are complementary to the distinctive quaternary structure of RapZ (Figure 2b). SLI and SLII make similar contacts to their corresponding protomers in RapZ, involving in many instances the same amino acid positions.

The RNA stem loops are primarily contacted by the CTD dimers of RapZ; SLI interacts with the CTD dimer formed by chains C and D, whilst SLII interacts with the CTD dimer formed by chains A and B (Figure 2b). For both of the stem loops the path of the RNA across the CTD dimer has approximately two-fold symmetry, and the conformation of the duplex changes from A-form to a highly underwound helix near the dyad symmetry axis (roughly between residue K170 of chains A and B, respectively, Figures 2e). The phosphate backbones of each of the two stem loops of GlmZ align with corresponding putative ligand-binding pockets formed by the CTDs of chains A/B and C/D, respectively. These latter pockets are composed of residues H190, T248, G249, H252 and R253 with C247 being in close vicinity, respectively (Figures 2e, 3a). Sulphate and malonate ions were previously seen at these positions in crystal structures of apo RapZ (Gonzalez et al., 2017). Several basic residues (K170, R184, K194, R196, R211, R238, K251) form patterns of positive charges on the CTD surface providing electrostatic complementarity to the path of the sRNA (Figure 2e). The pocket has shape and electrostatic complementarity to the path of one strand of the A-form RNA helix involving interactions with nucleotides A125 and A127 in SLII of GlmZ (Figure 2d, lower panel). The groove shape of the A-form helix is recognised by an exposed loop comprising RapZ residues 190-197. The sulphate-binding pocket has been proposed to also be the site for binding of glucosamine-6-phosphate (Gonzalez *et al*., 2017), which is supported by recent studies showing binding of the metabolite to the CTD of RapZ (Khan et al., 2020). The Walker A and B motifs located in the N-terminal domain of RapZ are not positioned to make contact with the RNA substrate, instead they appear to run along the opposite diagonal of the RapZ tetramer to the RNA (Figure 2b).

**Figure 3.**
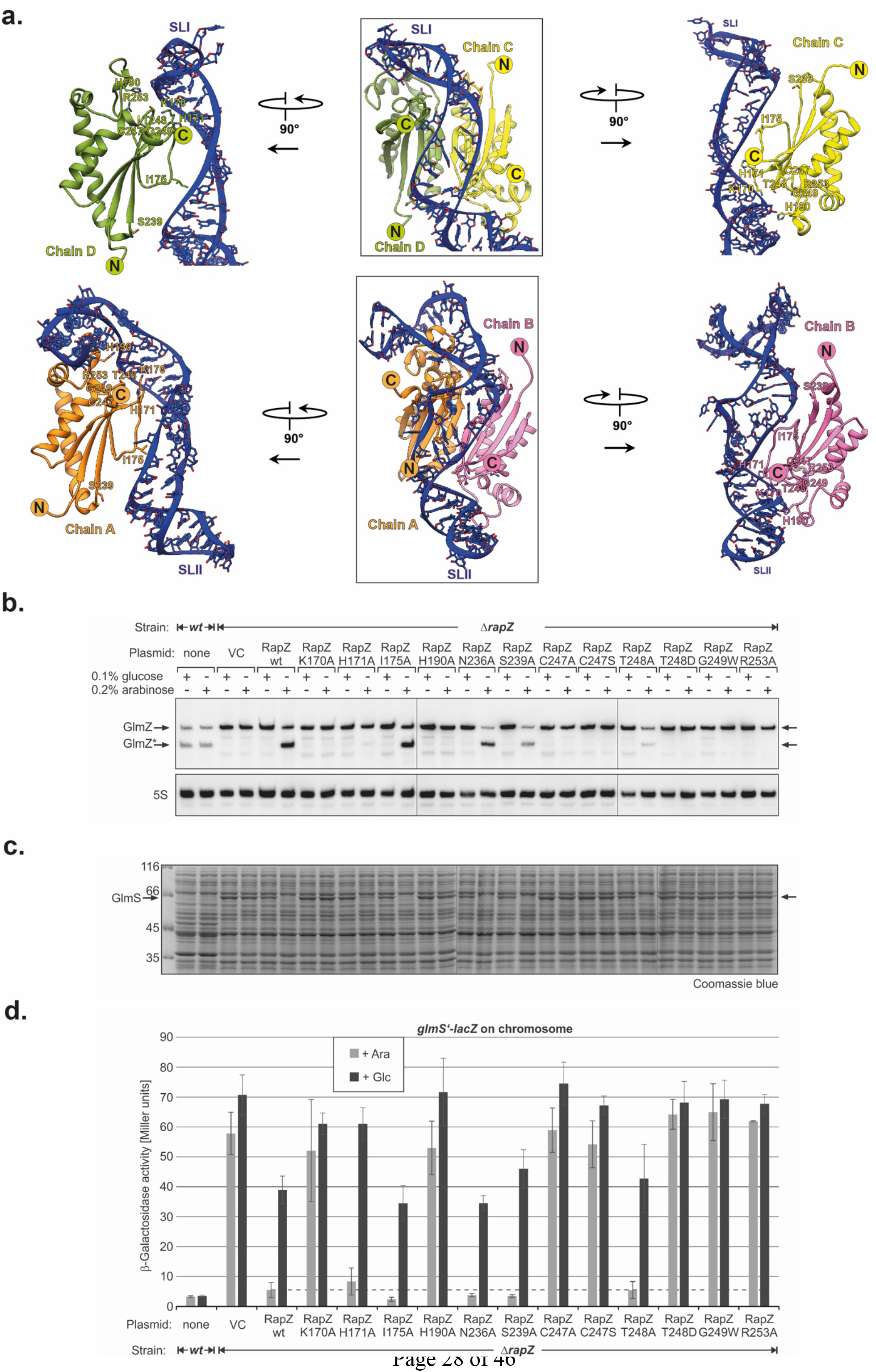
Importance of RapZ-GlmZ key contacts for regulatory activity of RapZ *in vivo*. (a) Interaction of the RapZ-CTD dimers with SLI (top) and SLII (bottom) of GlmZ. The indicated residues were substituted in RapZ and resulting variants were tested for regulatory activity *in vivo*. The *rapZ* mutants were expressed from plasmids under control of the *PBAD* promoter (induction with arabinose, repression by glucose) and tested for the ability to complement a strain lacking the chromosomal *rapZ* gene in **b.-d.** (b) Northern blotting experiment detecting GlmZ (top panel) and 5S rRNA (bottom panel; loading control) to assess the degree of GlmZ cleavage triggered by the RapZ variants. Processed GlmZ is indicated with an asterisk. (c) GlmS protein (MW = 66.9 kDa) levels in the same cultures as revealed by SDS-PAGE of total protein extracts followed by Coomassie blue staining. (d) β-Galactosidase activities produced by the same cultures and replicates (mean values and standard deviations are shown). Cells carry a *glmS’-lacZ* fusion on the chromosome, which requires base-pairing with sRNA GlmZ for high expression. Strains Z8 (*wild-type*, lanes/columns 1, 2) and Z28 (*ΔrapZ*, all other lanes/columns) were used. Strain Z28 carried plasmid pBAD33 (lanes/columns 3, 4; empty vector control = VC) or derivatives triggering synthesis of the indicated RapZ variants (cf. Table S2 for corresponding plasmids).

The RNase E cleavage site in GlmZ, which resides within the SSR located between SLII and SLIII, is not resolved in our cryoEM maps and is likely disordered. Thus, while interactions of GlmZ with RapZ present the GlmZ cleavage site to RNase E, it does not appear to pre-organise the substrate for enzyme attack. The binary complex is also likely to resemble the post-cleavage product, where only the SLI and SLII are bound by RapZ. Thus, processed GlmZ may interact with RapZ through contacts made by SLI and SLII as observed for full-length GlmZ. This explains earlier observations that accumulation of processed GlmZ counteracts cleavage through competition for RapZ, thereby providing feedback regulation (Durica-Mitic and Görke, 2019).

### RapZ:GlmZ contacts in the binary complex are crucial for regulatory activity in vivo

The residues of all four sulphate/malonate-binding pockets are observed in close vicinity to the phosphate backbone of GlmZ. The pockets of chains C and D huddle against the strands composing the A-form helix of SLI and the pockets in chains A/B are making contacts to SLII of GlmZ (Figures 2e, 3a). Hence, residues of these pockets should be key for proper orientation of the sRNA on the RapZ tetramer and its correct presentation to RNase E for cleavage. If so, their mutation is expected to interfere with cleavage of GlmZ by RNase E in the living cell, resulting in synthesis of GlmS at high levels (Kalamorz et al., 2007; Gonzalez et al., 2017). To test this, we changed four residues of the sulphate/malonate-binding pocket in RapZ (H190, T248, G249, R253) with disruptive substitutions. Additionally, we substituted nearby located residues K170, H171, I175, N236, S239 and C247, which might contribute to correct sRNA binding (Figure 3a). The *rapZ* mutants were expressed from plasmids under control of the arabinose-inducible *PBAD* promoter and analysed for complementation of an *E. coli* strain lacking the chromosomal *rapZ* copy. Western blot analysis confirmed synthesis of the RapZ variants upon induction with arabinose (Figure S6). Northern blot analysis confirmed that cleavage of GlmZ by RNase E is abolished in the *ΔrapZ* strain carrying the empty vector (VC, Figure 3b, lanes 3-4). The failure to degrade full-length GlmZ strongly activates *glmS’* translation as monitored by an ectopic *glmS’-lacZ* reporter fusion (Figure 3d), leading to accumulation of GlmS protein visible even in stained total protein extracts (Figure 3c, lanes 3-4). The presence of wild-type RapZ, either encoded endogenously or expressed from a plasmid, promotes processing of GlmZ and suppresses *glmS* expression as expected from previous work (Figures 3c, d, lanes 1-2, 6; Durica-Mitic et al., 2020).

Importantly, the substitutions H190A, G249W and R253A – all located in the sulphate/malonate-pocket - abolished the ability of RapZ to promote GlmZ cleavage and repress *glmS* (Figure 3b-d). RapZ remained unaffected when substituting T248 with Ala, but lost activity when replaced with an Asp, supporting our notion that electrostatic complementarity is important to interact with the negatively charged RNA backbone. This is further supported by inactivity of the K170A variant (Figure 3b-d), likely reflecting loss of a direct interaction with the nearby located RNA backbone (Figures 2e, 3a). Finally, Ala and Ser substitutions of C247, which is close to the sulphate/malonate-pocket (Figures 2e, 3a) abolish RapZ activity, albeit this was somewhat unexpected for the C247S variant. Perhaps, the -SH group fulfils a specialized function that could not be fulfilled by an -OH group. From the remaining variants, the Ala substitutions of I175, N236 and S239 had no effect and substitution of H171 lowered repression of *glmS* to some degree (Figure 3b-d).

Collectively, the results show that K170, C247 as well as the residues of the sulphate/malonate-pocket are essential for RapZ to promote cleavage of GlmZ *in vivo*. These observations are in agreement with the roles of these residues for presenting GlmZ correctly to RNase E, as predicted by the structure of the binary complex (Figure 2).

### Architecture of the RNaseE-NTD:RapZ:GlmZ complex

Extensive rounds of iterative 2D and 3D classifications revealed the presence of multiple states of the RNase E-NTD:RapZ:GlmZ complex, all containing a single RNase E-NTD tetramer but with varying occupancies of associated RapZ tetramers (Figure S7). However, an RNase E-NTD tetramer bound to one RapZ:GlmZ binary complex was common to all states, and we focussed on this fundamental unit to generate a structural model of the RNase E-NTD:RapZ:GlmZ ternary complex (Figure S8).

From our cryoEM map the quaternary organisation of the complex can be defined with confidence, as well as the general path of the RNA over the surface of the two proteins (Figures 4, S9 and Table S1). The binary complex of RapZ:GlmZ and the crystal structure of the tetrameric NTD of RNase E both fit well into the experimental map with small domain movements. Thus, the interaction of RapZ:GlmZ with RNase E does not involve conformational rearrangement of either RapZ or GlmZ. Previously published crystal structures of RNase E-NTD have revealed many degrees of freedom at both the quaternary and tertiary levels, enabling flexible accommodation of substrates (Callaghan et al., 2005; Koslover et al., 2008; Bandyra and Luisi, 2018; Bandyra et al., 2018). It may therefore not be surprising that in the ternary complex structure the 5′-sensor/S1 domains of the RNase E-NTD have adopted an opened conformation that can accommodate the binary complex – a configuration not previously observed but which may account for the capacity of RNase E to be guided to preferred cleavage sites by RNA structure (Figure S10).

**Figure 4.**
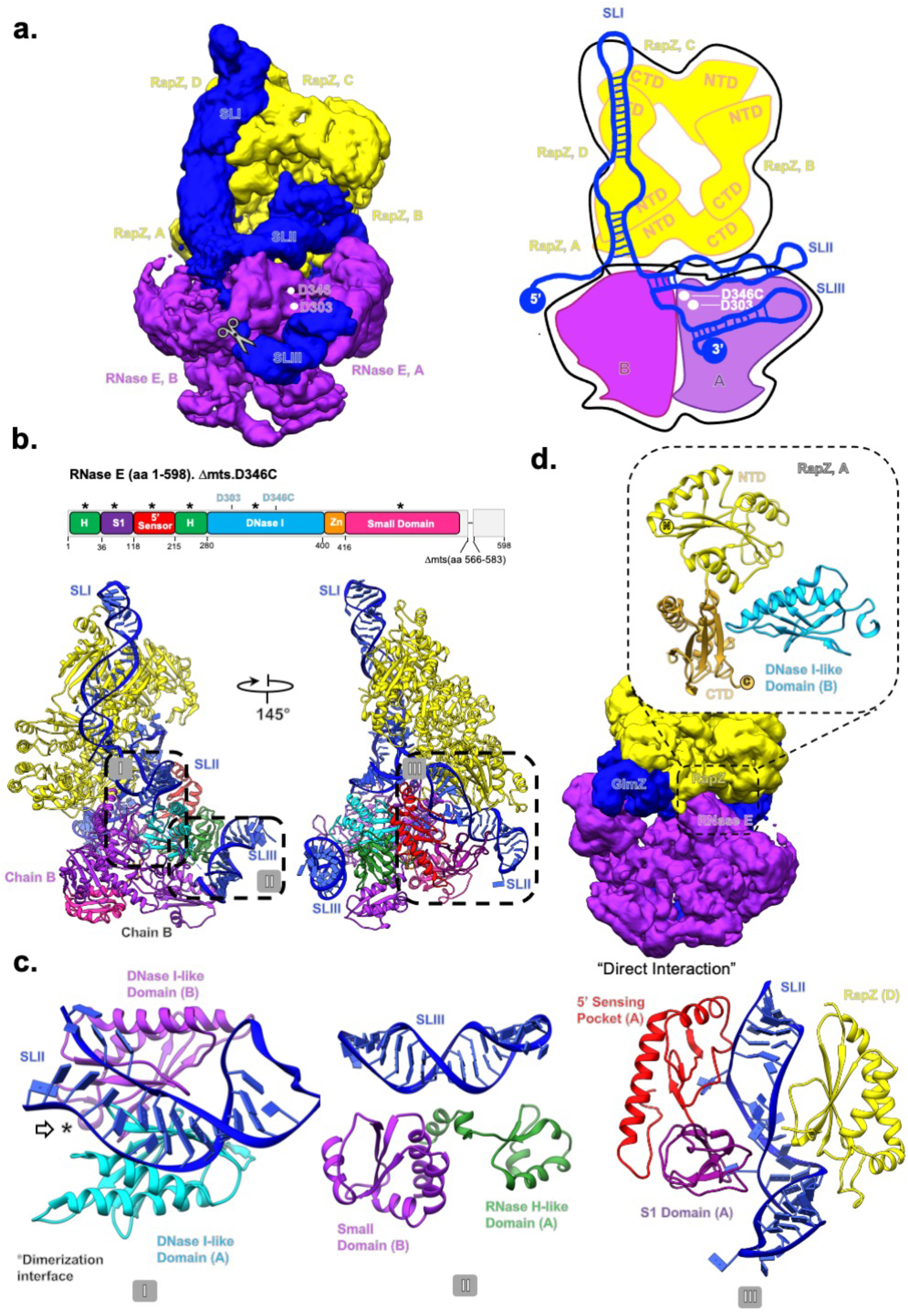
The RNaseE-NTD:GlmZ:RapZ ternary complex. (a) CryoEM map of the ternary complex, representing only the fundamental unit (depicted schematically in the cartoon in the right panel). (Figure S8 also provides a perspective of the fundamental unit with respect to the full complex.) (b) Model of the fundamental unit with the individual RNase E-NTD domains coloured according to the domain topology scheme. “Asterisks” on top of the schematic highlight the RNase E domains that interact with the binary RapZ-GlmZ complex. Boxes indicated with roman numerals I, II, and III indicate regions shown in expanded view in (c). (c) Detail views of the interactions between the RNase E-NTD and the sRNA: (c, left) Interaction of the RNase E DNase I-like domain with GlmZ-SLII, mediated via the dimerization interface of two DNase I domains from two nearby RNase E protomers. (c, middle) Interaction of the GlmZ-SLIII duplex with the RNase H-like domain of the cleaving protomer and the small domain of the nearby non-cleaving protomer. (c, right) Interaction between the 5′-sensing pocket and S1 domain of the non-cleaving protomer and GlmZ-SLII; this interaction facilitates interaction with the CTD of one of the RapZ protomers (chain B) using GlmZ as a bridge, further promoting formation of the overall complex. (d) Illustration of the direct interaction of RNase E with RapZ through a confined contact region.

### Recognition of RNA by RNase E catalytic domain ternary complex

The ternary complex assembly is stabilised predominantly through mutual binding of the RNase E-NTD and RapZ to GlmZ RNA (Figure 4). In particular, SLII of GlmZ, which was reported to be crucial for RNase E mediated cleavage (Göpel et al., 2016), appears to be a shared binding site for both RNase E and RapZ (Figure 4b-d). To gain insight into these interactions, we used the adenylate cyclase-based bacterial two-hybrid assay (BACTH). We reasoned that substitution of residues in RapZ that are required to bind SLII in a correct manner, should also affect the mutual contacts made by RNase E and thereby RapZ-RNase E interaction as measurable by BACTH in the *E. coli* cell (Durica-Mitic et al., 2020). Indeed, several substitutions including those located within the sulphate/malonate pocket decreased interaction fidelity significantly, with the H190A substitution yielding the strongest effect (Figure S11). These results support the idea that RapZ must present GlmZ correctly to facilitate mutual contacts by RNase E. However, removal of individual RapZ/GlmZ contacts is not sufficient to disrupt the ternary complex completely, likely reflecting remaining interactions (see also below).

The predicted structure of SLIII fits well into density present on the surface of the RNase E small domain and RNase H fold (Figures 4c and S10)). This binding site agrees with the earlier findings from an X-ray crystallographic structure of the RNase E-NTD in complex with the sRNA RprA, which indicated stem loop engagement on that surface (Bandyra et al., 2018). The small domain of RNase E is a divergent member of the KH domain family (Pereira and Lupas, 2018) and has a distinctive mode in RNA binding (Bandyra et al., 2018).

In a subset of particles, density continues from the base of the GlmZ SLIII structure in the direction of the active site in the DNase I sub-domain of RNase E. The location of the RNA in the active site is consistent with the position of the cleavage site in the SSR between SLII and SLIII in GlmZ (Göpel et al., 2016; Figures 2 and 4). It also agrees with the polarity of the phosphate backbone in the active site required to achieve the necessary geometry for cleavage (Callaghan et al., 2005). The S1 domain of RNase E encages the RNA in the active site, in agreement with earlier crystallographic studies (Callaghan et al., 2005). The 5′ sensor-S1 domain is comparatively disordered in relation to other regions within the map, and there is a gap in modelling the RNA in this region.

Continuing along the RNA in the 3′ to 5′ direction, there is clear density for SLII that engages the exposed surface of the DNase I domains from two nearby RNase E protomers (Figure 4c, left panel). This is the first observation of a duplex RNA interaction with this surface of RNase E; notably the SLII seems to interact *via* the dimerization interface of two DNase I domains. The duplex RNA continues and makes interactions with the CTD of the RapZ chain D, where it changes direction and then continues over the surface of the CTD of the RapZ chain C (Figure 4a). The duplex region contacted by RapZ chain A is also contacted by the 5′ sensor domain/S1 domain of RNase E, also revealing for the first time the interactions of this domain with a duplex RNA (Figure 4c, right panel; Figure S10). The interactions of the SLII with RNase E are predicted to be the key determinant for recognition of GlmZ as a substrate. Indeed, GlmY becomes a substrate for RapZ/RNase E *in vitro* if its SLII is replaced with that from GlmZ (Göpel et al., 2016). The base of SLII is near the active site, where the duplex region of SLI begins to continue over the surface of RapZ and interacts with the CTD of the RapZ chains A and B.

While the ternary complex of RNase E, GlmZ and RapZ is predominantly held together by protein-RNA interactions, there are also a few protein-protein interactions. Notably, the DNase I domain of RNase E chain B directly interacts with chain A of the RapZ tetramer (Figure 4d). The model reveals a small direct contact surface between RapZ and RNase E, formed by nestling of the DNase I domain between the N- and C-terminal domains of a proximal RapZ protomer. Here, RNase E segment 311-316 interacts with the surface of RapZ that includes residues Q273, N271, Y240, and T161. This complex interaction surface is consistent with results from mutagenesis screening performed in combination with bacterial two-hybrid assays (Durica-Mitic *et al*., 2020). The latter analysis showed that removal of a few residues from the RapZ C-terminus abolished interaction with RNase E and also indicated a critical role of residue L279 for direct interaction with RNase E. These observations are rationalised by the cryoEM model indicating that the RapZ C-terminus including residue L279 may support correct presentation of the strand that carries the contacting residues Q273 and N271.

## DISCUSSION

Removal of the seed pairing region from GlmZ by RNase E is a critical step in regulation of GlmS abundance and thereby for bacterial cell envelope biogenesis. Gram-negative Enterobacteria employ the adaptor protein RapZ to ensure tight control of this step to match the requirements of the cell during growth and stress response. Mechanistically, the question arises why cleavage of GlmZ relies on RapZ and cannot be catalysed by RNase E on its own. The structural data presented here reveal that RapZ presents the sRNA in a manner that aligns its SSR comprising the cleavage site into the RNase E active center, and that portions of GlmZ near the intersection of SLI and SLII interact with RNase E to pre-organise the channel, thereby providing access to the active site.

RapZ interacts with SLI and SLII of GlmZ, mainly through its CTDs, which form swapped dimers. Each CTD dimer within the RapZ tetramer binds one stem loop, which is achieved through complementarity in shape and electrostatic charge to the phosphodiester backbone of the sRNA (Figure 2). This arrangement also accounts for the observation that a CTD-dimer on its own is sufficient to bind the sRNAs with reasonable affinity (Gonzalez *et al*., 2017; Durica-Mitic et al., 2020). However, the CTD-dimer is not sufficient to mediate cleavage of GlmZ by RNase E, and the structure of the RapZ tetramer accounts for this observation. Both the N- and C-terminal domains are required to form the quaternary structure that is needed to bind both RNA stem loops and present the single-stranded cleavage site appropriately to RNase E. The kinase-like N-terminal domain of RapZ (NTD) makes only a few interactions with the RNA, and the path of the RNA does not encounter the Walker A or B motifs (Figure 2b). It is possible that this domain could act as an allosteric switch, whereby the binding of an as yet unknown ligand triggers quaternary structural changes that affect RapZ functions. Several positively charged residues in the CTD provide electrostatic complementarity to the path of the sRNA (Figure 2e) and residues from putative metabolite pockets make contacts to both SLs of GlmZ (Figures 2, 3a). Substitution of these residues in RapZ abrogates cleavage of GlmZ in the cell (Figure 3b), confirming their importance for presenting GlmZ correctly to RNase E. Moreover, they decrease RapZ/RNase E interaction fidelity (Figure S11), suggesting that correct binding of the sRNA by RapZ is a prerequisite for subsequent recognition by RNase E to form the ternary cleavage complex. The far C-terminal tail of RapZ is in particular enriched for basic residues and is known to also contribute to RNA binding (Göpel et al., 2013; Durica-Mitic *et al*., 2020). The cryoEM maps of the binary RapZ:GlmZ complex indicate that this region comprising residues 281-284 is generally disordered. Nonetheless, in all four RapZ subunits the far C-terminus is in close proximity to GlmZ and likely forms distributed interactions with the phosphodiester backbone. In conclusion, the structural features revealed here perfectly rationalise the findings of earlier studies indicating that the RapZ-CTD binds the sRNA, but that the tetrameric structure is indispensable (Gonzalez et al., 2017; Durica-Mitic *et al*., 2020), because it engages two CTD-dimers to ensure binding of both SLs, which allows GlmZ to be presented to RNase E .

Inside the cell, cleavage of GlmZ by RNase E is counteracted by the homologous decoy sRNA GlmY, which accumulates when GlcN6P synthesis is required (Figure 1; Khan et al., 2020; Göpel et al., 2013). Based on the predicted fold, GlmY is likely to form a complex with RapZ that is structurally similar to the RapZ:GlmZ binary complex. Notably, the species accumulating *in vivo* refers to a processed GlmY variant consisting only of SLI and SLII (Reichenbach et al., 2008; Urban and Vogel, 2008). SLIII does not contribute to recognition of the sRNA by RapZ (Figure 2), which explains why short GlmY is nonetheless capable to bind and sequester RapZ efficiently. The model also explains why the post-cleavage product, *i.e.* GlmZ lacking SLIII, can still inhibit GlmZ cleavage in a negative feed-back loop, when accumulating above a threshold (Durica-Mitic and Görke, 2019). RapZ could in principle bind many RNA species in the cell, but analysis of RNA copurifying with RapZ in pull-down experiments identified only GlmY and GlmZ as substrates (Göpel et al., 2013). This high specificity arises from the requirement for binding two distinct RNA stem-loops through a unique pattern of complementary shape and charge in RapZ (Figure 2).

*In vivo*, the adapter function of RapZ is restricted to presenting GlmZ to RNase E for cleavage, at least under the growth conditions analysed so far (Durica-Mitic *et al*., 2020). This specificity is not simply a consequence of the RNA-binding specificity of RapZ. For instance, although full-length GlmY is readily bound by RapZ, it is not cleaved in the GlmY:RapZ binary complex by RNase E in cleavage assays *in vitro* (Göpel et al., 2016). However, GlmY becomes cleavable when its SLII is swapped for its counterpart from GlmZ, suggesting that this structure is decisive for recognition by RNase E. SLII of GlmZ contains unique features such as the lateral bulge in the SLII (Figure 2a) that is part of an underwound helix contacted by both RapZ and RNase E in the encounter complex (Figures 2d, 4b). The somewhat different SLII of GlmY is likely unable to mediate this mutual binding to RapZ and to the 5′ sensor-S1 domain of RNase E. Additionally, the SSR following SLII requires alignment in correct polarity with the catalytic site of RNase E to get cleaved (Figures 4a, 4b). This task could be facilitated by the downstream terminator stem loop structure (i.e. SLIII) that engages the RNase H/KH-like small domain in RNase E. This recognition may be guided by RNA secondary structure and less by sequence, considering that RNase E binds the phosphodiester backbone of an A-form helix segment in SLIII (Figure 4c, middle panel). Previous structural studies on a distinct sRNA/RNase E complex also indicated stem loop engagement on that surface (Bandyra et al., 2018) and molecular biology studies showed that SLIII of GmZ can be replaced with non-cognate terminators without affecting cleavage (Göpel et al., 2016). Therefore, it appears reasonable that the RNase H/KH-like small domain provides a general RNA docking site that could also play a role for formation of encounter complexes with other RNA chaperones (see below). Finally, the mode of recognition of GlmZ by RapZ and RNase E suggests key features to engineer cleavage sites that could be used to trigger *in vivo* genesis of defined RNA species from precursors (Göpel et al., 2016).

Interestingly, the phosphate groups of the RNA backbone occupy positions in RapZ that were previously observed to bind sulphate or malonate ions in the crystal structure of apo-RapZ, suggesting that this pocket could be the binding site for a charged metabolite such as GlcN6P (Figure 2e, 3a; (Gonzalez et al., 2017)). GlcN6P is known to bind to the RapZ-CTD at an as yet uncharacterized site, thereby interfering with sRNA binding and stimulation of QseE/QseF activity (Khan et al., 2020). Strikingly, the structure of the binary complex predicts mutually exclusive access of the metabolite and the sRNA to the sulphate/malonate pocket, reinforcing the idea that GlcN6P binds here. It has been noted earlier that the fold of the C-terminal domain of RapZ closely resembles the fructose-6-phosphate binding domain of the metabolic enzyme phosphofructokinase (Gonzalez *et al*., 2017); Figure 2b)). Metabolic enzymes have been identified in moonlighting functions as RNA binding proteins with roles in post-transcriptional regulation (Beckmann *et al*., 2015; Marondedze *et al*., 2016). Our observations hint as to how these enzymes might recruit RNA species through reallocation of metabolite-binding sites or be repurposed in the course of evolution for this role, adding to the discussion of possible relationships between metabolic enzymes and RNA-binding proteins (Hentze *et al*., 2018; Balcerak et al., 2019; Corley et al., 2020).

For the ternary RNaseE-NTD:RapZ:GlmZ complex, the predominant stoichiometry we observed was one RNase E-NTD tetramer, two RapZ tetramers and two GlmZ molecules. However, the binding of the RapZ:GlmZ binary assemblies to the RNase E-NTD can occur in *cis* or *trans* conformations (with the binary complexes engaging on the same or opposite faces of the RNase E-NTD) (Figure S7). These higher order complexes may result from saturation of possible binding sites and are not anticipated to be required mechanistically. However, they do illustrate the flexibility of the RNase E tetramer and the potential for the ribonuclease to undergo conformational adaptation to accommodate larger substrates through multiple contacts mediated by distant sites. This mode of binding could play a role in the capacity of RNase E to migrate on long RNA substrates (Richards and Belasco, 2019, 2021).

The structures studied here may also provide a clue as to the mechanism for other effector complexes known to engage RNase E. For instance, the regulatory protein CsrD, which is required for specific turnover of the sRNAs CsrB and CsrC by RNase E (Vakulskas *et al*., 2016; Potts *et al*., 2018), may present those RNAs for structure-based recognition in analogy to the RapZ/GlmZ complex studied here. A similar scenario could also apply to recently identified RNA-binding proteins in non-model bacteria such as CcaF1, which modulates degradation of selected transcripts by RNase E in *Rhodobacter sphaeroides* (Grützner et al., 2021), or to the emerging class of KH proteins that may act as sRNA chaperones in Gram-positive bacteria (Olejniczak et al., 2021). It can even be envisaged that the global RNA chaperone Hfq may form encounter complexes analogous to RapZ to facilitate degradation of base-paired sRNA/target RNAs by RNase E (Figure 5a, right panel). In this model, the sRNA/target duplex would be recognized by both, Hfq and RNase E, reminiscent of the role of GlmZ-SLII in the current encounter complex. The sRNA 3’ poly (U) end may stay bound on Hfq while the newly identified duplex binding site on RNase E could bind the structured target RNA to accommodate the cleavable site in the catalytic domain (Figure 2b). The new duplex binding region in RNase E may even have general implications for how binding of other structured RNA substrates may occur. For instance, processing of polycistronic tRNAs may involve capture complexes in which a duplex region of the tRNA is engaged on RNase E to generate a defined cleavage site (Kime et al., 2014). The 3’ located tRNA may be docked onto the RNase H/small domain surface while engaging the preceding tRNA copy on the exposed S1/5’sensor domain. The results presented here provide insight into the mechanism of substrate preferences for RNase E and into the role of RNA chaperones to guide the activity of this key enzyme of RNA metabolism.

**Figure 5.**
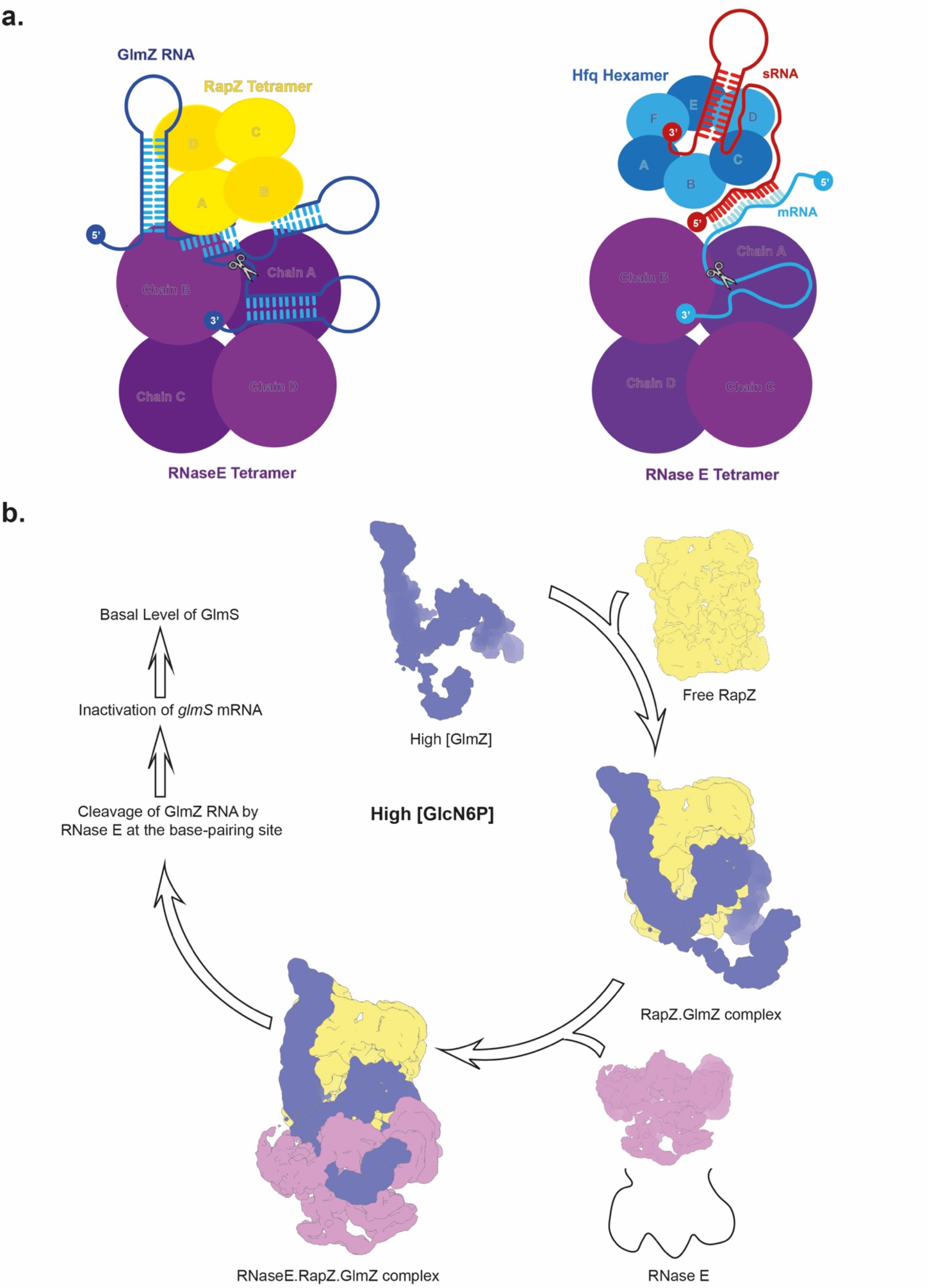
Proposed models for presentation of RNA-duplexes by RNA chaperones to RNase E. (a) The organisation of the RNase E-NTD:GlmZ:RapZ complex (left) suggests an analogous mode for presentation of complex double-stranded RNA substrates by other RNA-chaperones such as Hfq (right). A hypothetical encounter complex is shown illustrating how Hfq may deliver base-paired small RNAs (red) and RNA targets (blue) to RNase E for cleavage. An mRNA is indicated in the cartoon, but targets could also include non-coding RNAs such as GcvB. (b) Model illustrating the mechanism of GlmZ cleavage according to the current work. As a prerequisite for cleavage, GlmZ must be bound by the tetrameric adapter protein RapZ. Binding occurs through recognition of SLI and SLII in GlmZ by the CTD-dimers of RapZ, whereas SLIII has no role. This mode of recognition explains earlier observations that the processed variants of GlmY as well as GlmZ carrying SLI and SLII only, compete successfully with full-length GlmZ for binding RapZ (Göpel et al., 2013; Durica-Mitic and Görke, 2019). Correct presentation of GlmZ by RapZ allows joining of RNase E to form the ternary encounter complex, which is stabilized through mutual interaction of the proteins with SLII of GlmZ, binding of the GlmZ-SLIII to the RNase H-like/small domain surface on RNase E and direct RapZ-RNase E contacts. This assembly positions the SSR of GlmZ in the RNase E active site, allowing for cleavage. The resulting GlmZ molecule lacks the base-pairing site and is not capable to activate *glmS* translation.

## Materials and Methods

### Construction of rapZ mutants with single codon exchanges

Single codon exchanges were introduced into *rapZ* using the combined chain reaction (CCR) method (Bi & Stambrook, 1998) and resulting mutants were placed on plasmid pBAD33 under control of the arabinose-inducible *PBAD* promoter. Briefly, 5’-phosphorylated oligonucleotides carrying the desired nucleotide exchanges were incorporated into *rapZ* by thermostable ampligase (Biozym Scientific GmbH) during amplification by PCR using the forward/reverse primers BG1049/BG397. DNA fragments were digested and inserted between SacI/XbaI restriction sites on plasmid pBAD33. Plasmid constructs for the bacterial adenylate cyclase-based two-hybrid assay (BACTH) were obtained by CCR reactions using the forward/reverse primers BG637/BG639. DNA-fragments were cloned between the XbaI-KpnI sites on plasmid pKT25 to generate corresponding *T25-rapZ* fusion genes. The resulting plasmids and corresponding 5’-phosphorylated oligonucleotides are listed in Table S2. Oligonucleotide sequences are indicated in Table S3. Mutations were verified by DNA sequencing.

### Analysis of regulatory activity of RapZ variants and their interaction with RNase E in vivo

Plasmid pBAD33 and its various derivatives encoding the RapZ variants were introduced into the *ΔrapZ* mutant strain Z28, respectively, and resulting transformants were grown overnight in LB containing 15 μg/ml chloramphenicol and supplemented either with 0.1% glucose or with 0.2% L-arabinose. For comparison, the isogenic *wild-type* strain Z8 (*rapZ^+^*) was included, but chloramphenicol selection was omitted in this case. On the next day, cells were inoculated to an OD600 = 0.1 into fresh medium and grown until cultures reached an OD600 ∼ 0.5-0.8. Subsequently, samples were harvested for isolation of total RNA, total protein and determination of β-galactosidase activities, respectively. β-Galactosidase activities were measured as described (Miller, 1972). Presented values represent the average of at least three measurements using at least two independently transformed cell lines. For sodium dodecyl sulphate-polyacrylamide gel electrophoresis (SDS-PAGE) analysis, cells of each culture corresponding to 0.05 OD600 units were dissolved in SDS sample buffer and separated on 10% SDS polyacrylamide gels, which were subsequently stained with Coomassie brilliant blue R-250 or subjected to Western blotting using a polyclonal RapZ antiserum (Durica-Mitic and Görke, 2019). Extraction of total RNA and Northern blotting was performed as described previously (Durica-Mitic *et al*, 2020), unless 2.5 μg total RNA of each sample were separated on denaturing gels containing 7 M urea, 8% acrylamide and 1× TBE. RNase E/RapZ interactions were analysed using the bacterial adenylate cyclase-based two-hybrid system (BACTH) according to an established protocol (Durica-Mitic et al., 2020). Briefly, transformants of *E. coli* reporter strain BTH101 carrying the desired combinations of pUT18C-and pKT25-type plasmids were grown overnight to early stationary phase in LB (100 μg/ml ampicillin, 30 μg/ml kanamycin, 1 mM isopropyl-β-thiogalactopyranoside (IPTG) and the β-galactosidase activities were determined.

### RNase E NTD expression and purification

Purification of RNase E encompassing residues 1 to 598 (NTD) without the membrane targeting sequences (Δ567-583) and with the active-site residue Asp^346^ mutated to cysteine (D346C), wild-type RNase E-NTD (aa 1-529) followed protocols for shorter versions of the catalytic domain (Callaghan *et al*., 2005; Bandyra et al., 2018). *Escherichia coli* strain BL21(DE3) was transfected with pET16 expression vector overproducing RNase E variants with an *N*-terminal his6-tag (kindly provided by A.J. Carpousis). Cultures were grown in 2xTY media supplemented with 100 µg/mL carbenicillin at 37°C, using dimpled flasks for optimal aeration, in an orbital shaker set at 160 rpm. The culture was induced between 0.5 to 0.6 OD600 by adding 1 mM IPTG and harvested after 3 hours of incubation by centrifugation at 5020 g and 4° C for 30 minutes. Cell pellets were stored as suspension in nickel-column buffer A (20 mM Tris pH 7.9, 500 mM NaCl, 5 mM imidazole, 1 mM MgCl2) at -80°C until further use. Once thawed, the cell culture suspension was supplemented with DNase I and EDTA-free protease inhibitor cocktail tablet (Roche) and lysed by passing through an EmulsiFlex-05 cell disruptor (Avestin) for 2-3 times at 10-15 kbar pressure. The lysate was clarified by centrifugation at 35000 g for 30 minutes at 4° C and the supernatant was passed through a 0.45 µ membrane filter before loading onto a pre-equilibrated HiTrap immobilized metal ion affinity (IMAC) column (HiTrap IMAC FF, GE Healthcare). The column was washed extensively with wash buffer (20 mM Tris pH 7.9, 500 mM NaCl, 100 mM imidazole, 1 mM MgCl2), and RNase E eluted by a gradient of elution buffer (20 mM Tris pH 7.9, 500 mM NaCl, 500 mM imidazole, 1 mM MgCl2). Protein quality was assessed by SDS-PAGE and fractions containing RNase E were pooled and loaded onto a butyl sepharose column (GE Healthcare) equilibrated in high-salt buffer (50 mM Tris pH 7.5, 50 mM NaCl, 25 mM KCl, 1 M (NH)2SO4) to eliminate co-purifying nucleic acids. A low-salt buffer (50 mM Tris pH 7.5, 50 mM NaCl, 25 mM KCl, 5% v/v glycerol) was used for the elution. Based on purity as determined by SDS-PAGE, fractions were pooled, concentrated using a 50 kDa MWCO concentrator, and loaded onto a size-exclusion column (Superdex^TM^ 200 Increase 10/300, GE Healthcare) equilibrated in storage buffer (20 mM HEPES pH 7.0, 500 mM NaCl, 10 mM MgCl2, 0.5 mM TCEP, 0.5 mM ethylenediaminetetraacetic acid (EDTA), 5% v/v glycerol). The best fractions were flash frozen in liquid nitrogen and stored at -80 °C until further use.

### RapZ expression and purification

Full-length RapZ protein was expressed and purified by following a previously reported procedure (Gonzalez *et al*., 2017). Briefly, *Escherichia coli* Rosetta cells were transformed with plasmid pMCSG7 carrying the *gene* encoding full-length RapZ with a Strep-tag cleavable by TEV protease. Bacterial cultures were grown in LB media supplemented with 1% (w/v) glucose, 30 µg/mL chloramphenicol, and 100 µg/mL carbenicillin at 37°C in dimpled flasks for optimal aeration and in an orbital shaker set at 220 rpm. The cultures were induced at OD600 ∼0.8 by adding 1 mM IPTG, grown at 18°C for another 1 hour and finally harvested by centrifugation at 5020 g and 4° C for 30 minutes. Cell pellets were stored in lysis buffer (50 mM Tris pH 8.5, 500 mM KCl, 1 mM EDTA, 5 mM β-mecaptoethanol) at -80°C until further use. Once thawed, the cell culture suspension was supplemented with DNase I and EDTA-free protease inhibitor cocktail tablet (Roche) and cells were lysed by passing through an EmulsiFlex-05 cell disruptor (Avestin) for 2-3 times at 10-15 kbar pressure. The lysate was clarified by centrifugation at 35000 g for 30 minutes at 4° C and the supernatant was passed through a 0.45 µm membrane filter before loading onto a pre-equilibrated Strep Trap HP column (GE Healthcare) equilibrated previously with Strep Trap buffer A (50 mM Tris pH 8.5, 500 mM KCl, 1 mM EDTA, 5 mM β-mecaptoethanol). The column was washed extensively with buffer A before protein was eluted with buffer B (50 mM Tris pH 8.5, 500 mM KCl, 1 mM EDTA, 5 mM β-mercaptoethanol, and 2.5 mM desthiobiotin). Protein quality was judged by SDS-PAGE and fractions containing RapZ were pooled. A final purification step was carried out by size-exclusion chromatography using a Superdex 200 Increase 10/300 (GE Healthcare) column equilibrated with storage buffer (50 mM Tris pH 7.5, 100 mM NaCl, 50 mM KCl, 2 mM dithiothreitol). The fractions enriched in RapZ, as judged by SDS-PAGE, were flash frozen in liquid nitrogen and stored at -80 °C until further use.

### RNA preparation by in vitro transcription

RNAs were produced by *in vitro* transcription. Templates were prepared by polymerase chain reaction (see Table S3 for oligonucleotides) and GlmZ and GlmZ-Pro RNA were generated using T7 RNA polymerase at 37°C, followed by treating the reaction mixture with TURBO DNase for 15-20 minutes at 37°C to eliminate the DNA template. Synthesized GlmZ RNAs were purified on 6% 7.5 M urea polyacrylamide gel (National Diagnostics). The bands were visualized under a portable UV lamp at 254 nm wavelength and excised. Finally, the RNA was purified from the excised gel by overnight electroelution at 4° C and 100V (EluTrap, Whatman). For all RNAs, purity was checked by 8% urea-PAGE gel electrophoresis and SYBR gold RNA dye (Thermo Fisher) was used to visualize RNA (Figure S2d).

### Cryo-EM grid preparation

#### RNaseE-NTD:RapZ:GlmZ Complex

Purified RNase E-NTD, full-length RapZ, and full-length GlmZ RNA were incubated in a ratio of 5:25:10 µM in reaction buffer (25 mM HEPES pH 7.5, 300 KCl, and 1 mM MgCl2) supplemented with 0.01% glutaraldehyde at RT for 20 mins followed by at 30 °C for 10 minutes. The cross-linking reaction was quenched by adding 50 mM Tris pH 7.5. Samples were run on to a Superose 6 size-exclusion column equilibrated with the reaction buffer. Fractions showing evidence for protein elution (judged by SDS-PAGE) with high 260/280 ratio were pooled, concentrated to 1 mg/mL and applied to Quantifoil Cu 300 1.2/1.3 grids coated with graphene oxide (GO) (Palovcak et al. 2018; Russo and Passmore, 2014). Briefly, grids were glow-discharged on the darker carbon side in a PELCO easiGLOW glow discharge unit equipped with an oil pump (TED PELLA Inc., USA) using the following conditions: 15 mA, 0.28 mBar, 2 minutes. GO solution (2 mg/mL dispersion in water, Sigma-Aldrich product code: 763705) was 10x diluted using ultrapure water (ddH2O) and centrifuged at 300 x g for 30 secs to remove insoluble GO flakes. The supernatant was further diluted another 10x to make a 0.02 mg/mL GO solution. On the glow-discharged side of the Quantifoil grids, 1 µL of 0.02 mg/mL GO solution was applied and waited until the water evaporated. GO-coated grids were left at room temperature for at least 12-16 hrs before use to prepare EM specimens.

### Cryo-EM data acquisition and processing

Cryo-EM data were collected on a Titan Krios G3 in the Department of Biochemistry, University of Cambridge, with parameters given in Table S1. 2,445 movies were collected in accurate hole centering mode using EPU software (Thermo Fisher). CTF correction, motion correction, and particle picking were performed using (Tegunov and Cramer, 2019). 286172 particles picked by boxnet2 masked neural network model in Warp were imported to CryoSPARC (Punjani *et al*., 2017) for all subsequent processing. These particles were initially subjected to two-dimensional (2D) classification and 125,949 particle selected from these classes were used to generate initial *ab initio* 3D volumes representing RapZ:RNase E-NTD assemblies. The remaining 160,223 particles were also used to generate several ab initio 3D volumes to represent particles that do not contain RapZ:RNase E-NTD. Particles corresponding to different classes were selected and optimised through multiple iterative rounds of heterogeneous refinement as implemented in CryoSPARC. This process initially used the entire population of picked particles and all initial 3D volumes, and through the iterative process particles not representing RapZ assemblies were discarded from the set used for reconstructions. Particles representing the ternary complex of RapZ:RNase E-NTD:GlmZ were subjected to particle subtraction to remove the variable part of the molecule (within a user defined masked area). The best models were then further refined using homogenous refinement and finally non-uniform refinement in CryoSPARC. The classification process is summarised schematically in Figure S3. The final reconstructions obtained had overall resolutions (Table S1), which were calculated by Fourier shell correlation at 0.143 cut-off (Figures S6, S9).

### Structure refinement and model building

Crystal structures of full-length RapZ (PDB:5O5O) (Gonzalez *et al*., 2017) and RNase E-NTD (residues 1-510, PDBs: 2C0B, 6G63) (Callaghan *et al*., 2005; Bandyra, Wandzik and Luisi, 2018) were initially placed into the cryo-EM map. Further model building in COOT (Emsley and Cowtan, 2004) was followed by iterative cycles of refinement and density improvement in PHENIX (Afonine *et al*., 2018; Terwilliger *et al*., 2020). The RNA stem loop structures were predicted using the RNAfold web server (Lorenz et al. 2011) and subsequently subjected to the SimRNA server to generate 3D structures (Magnus *et al*., 2016, Boniecki *et al*., 2015). The structures were used as references to generate restraints for using PROSMART, and the models where docked into the cryoEM map and manually adjusted before local refinement.

## Acknowledgements

This work is supported by a Wellcome Trust investigator award to BFL (200873/Z/16/Z). Contributions from the Görke laboratory were supported by grant P32410-B of the “Austrian Science Fund” (FWF). We thank AJ Carpousis for the expression vector expressing RNase E-NTD with active site mutation. We thank Kasia Bandyra, Kotryna Bloznelyte, Modestas Matusevicius, Giulia Paris and Tom Dendooven for helpful discussions and Svetlana Durica-Mitic for help with plasmid constructions.

## Conflict of interests

The authors declare that they have no conflict of interest.

---

**Table S1.** Cryo EM data collection and refinement statistics

|  | Ternary | Binary |
| --- | --- | --- |
| **Data collection and processing** |  |  |
| Microscope | Titan Krios G3 | Titan Krios G3 |
| Detector | Gatan K3 | Gatan K3 |
| Magnification | 130,000 | 130,000 |
| Energy filter slit width (eV) | 20 | 20 |
| Voltage (kV) | 300 | 300 |
| Flux on detector (e/pix/sec) | 15.20 | 15.20 |
| Electron exposure on sample (e-/Å<sup>2</sup>) | 46.84 | 46.84 |
| Target defocus range (µm) | 1.2-2.6 | 1.2-2.6 |
| Calibrated pixel size (Å) | 0.652 | 0.652 |
| Symmetry imposed | C1 | C1 |
| Extraction box size (pixels) | 460 | 460 |
| Initial particle images (no.) | 286,172 | 286,172 |
| Final particle images (no.) | 33,595 | 27,551 |
| **Refinement** |  |  |
| Map resolution at FSC=0.143 (Å)* | 3.99 | 4.28 |
| Model composition |  |  |
| Non-hydrogen atoms | 19392 | 10848 |
| Protein residues | 2145 | 1126 |
| Nucleotides | 174 | 122 |
| B factor (Å<sup>2</sup>) |  |  |
| Protein | 239.82 | 250.07 |
| RNA | 281.69 | 256.37 |
| Bonds (RMSD) |  |  |
| Lengths (Å) | 0.013(33) | 0.014(17) |
| Angles (°) | 2.977(689) | 2.955(454) |
| Validation |  |  |
| Molprobity score | 2.45 | 2.57 |
| Clashscore | 19 | 20 |
| Poor rotamers (%) | 0.0 | 0.4 |
| Ramachandran plot |  |  |
| Favored (%) | 88.65 | 91.94 |
| Allowed (%) | 11.35 | 7.96 |
| Disallowed (%) | 0.0 | 0.1 |

**Table S2.** Strains and plasmids used in this study.

| Name | Relevant structure/genotype | Reference/construction<sup>a</sup> |
| --- | --- | --- |
| *E. coli* K-12 strains: |  |  |
| BTH101 | *F<sup>-</sup> cya-99 araD139 galE15 galK16 rpsL1 (Str<sup>R</sup>) hsdR2 mcrA1 mcrB1* | Karimova et al., 1998 |
| Z8 | CSH50 $\Delta$(*pho-bgl*)201 $\Delta$(*lac-pro*) *ara thi* ΔattB::[*aadA*, *glmS'*-*lacZ*] | (Kalamorz et al, 2007) |
| Z28 | as Z8, but Δ*rapZ* | (Kalamorz et al., 2007) |
| Plasmids: |  |  |
| pBAD33 | *P<sub>Ara</sub>*, MCS 2, *cat*, ori p15A | (Guzman et al, 1995) |
| pBGG61 | *rapZ* (-17 to +855) under *P<sub>Ara</sub>*-control in pBAD33 | (Göpel et al, 2013) |
| pBGG348 | encodes T25-RapZ in pKT25 | Göpel et al., 2013 |
| pBGG443 | as pBGG61, but encodes RapZ with Lys170Ala substitution | this work, using [P]-oligo BG1235 |
| pBGG449 | encodes T25-RapZ-Ser239Ala in pKT25 | this work, using [P]-oligo BG1350 |
| pBGG453 | as pBGG61, but encodes RapZ with Ser239Ala substitution | this work, using [P]-oligo BG1350 |
| pBGG457 | encodes T25-RapZ-Arg253Ala in pKT25 | this work, using [P]-oligo BG1369 |
| pBGG461 | as pBGG61, but encodes RapZ with Arg253Ala substitution | this work, using [P]-oligo BG1369 |
| pKT25 | *P<sub>lac</sub>::cyaA* [1-224] (T25), MCS, *neo*, ori p15A | Karimova et al., 1998 |
| pLQ18 | encodes T25-RapZ-His171Ala in pKT25 | this work, using [P]-oligo BG2071 |
| pLQ19 | encodes T25-RapZ-Ile175Ala in pKT25 | this work, using [P]-oligo BG2072 |
| pLQ20 | encodes T25-RapZ-Cys247Ser in pKT25 | this work, using [P]-oligo BG1731 |
| pLQ21 | encodes T25-RapZ-Thr248Asp in pKT25 | this work, using [P]-oligo BG2073 |
| pLQ22 | as pBGG61, but encodes RapZ with His171Ala substitution | this work, using [P]-oligo BG2071 |
| pLQ23 | as pBGG61, but encodes RapZ with Ile175Ala substitution | this work, using [P]-oligo BG2072 |
| pLQ24 | as pBGG61, but encodes RapZ with Cys247Ser substitution | this work, using [P]-oligo BG1731 |
| pLQ25 | as pBGG61, but encodes RapZ with Thr248Asp substitution | this work, using [P]-oligo BG2073 |
| pLQ33 | as pBGG61, but encodes RapZ with Asn236Ala substitution | this work, using [P]-oligo BG2115 |
| pLQ34 | encodes T25-RapZ-Asn236Ala in pKT25 | this work, using [P]-oligo BG2115 |
| pMCSG7 | encodes RapZ with a Strep-tag cleavable by TEV protease | Gonzalez et al., 2017 |
| pSD107 | as pBGG61, but encodes RapZ with His190Ala substitution | this work, using [P]-oligo BG1590 |
| pSD108 | as pBGG61, but encodes RapZ with Thr248Ala substitution | this work, using [P]-oligo BG1591 |
| pSD109 | as pBGG61, but encodes RapZ with Gly249Trp substitution | this work, using [P]-oligo BG1592 |
| pSD121 | encodes T25-RapZ-His190Ala in pKT25 | this work, using [P]-oligo BG1590 |
| pSD122 | encodes T25-RapZ-Thr248Ala in pKT25 | this work, using [P]-oligo |
|  |  | BG1591 |
| pSD123 | encodes T25-RapZ-Gly249Trp in pKT25 | this work, using [P]-oligo BG1592 |
| pSD130 | as pBGG61, but encodes RapZ with Cys247Ala substitution | this work, using [P]-oligo BG1480 |
| pSD139 | encodes T25-RapZ-Cys247Ala in pKT25 | this work, using [P]-oligo BG1480 |
| pUT18C | *P<sub>lac</sub>::cyaA* [225-399] (T18), MCS, *bla*, *ori* ColEI | Karimova et al., 1998 |
| pYG97 | encodes T18-RNase E (aa 1-597) fusion in pUT18C | Göpel et al., 2013 |

<sup>a</sup> [P]-oligo: 5' phosphorylated oligonucleotide for introduction of corresponding codon exchange into *rapZ*.

**Table S3.**
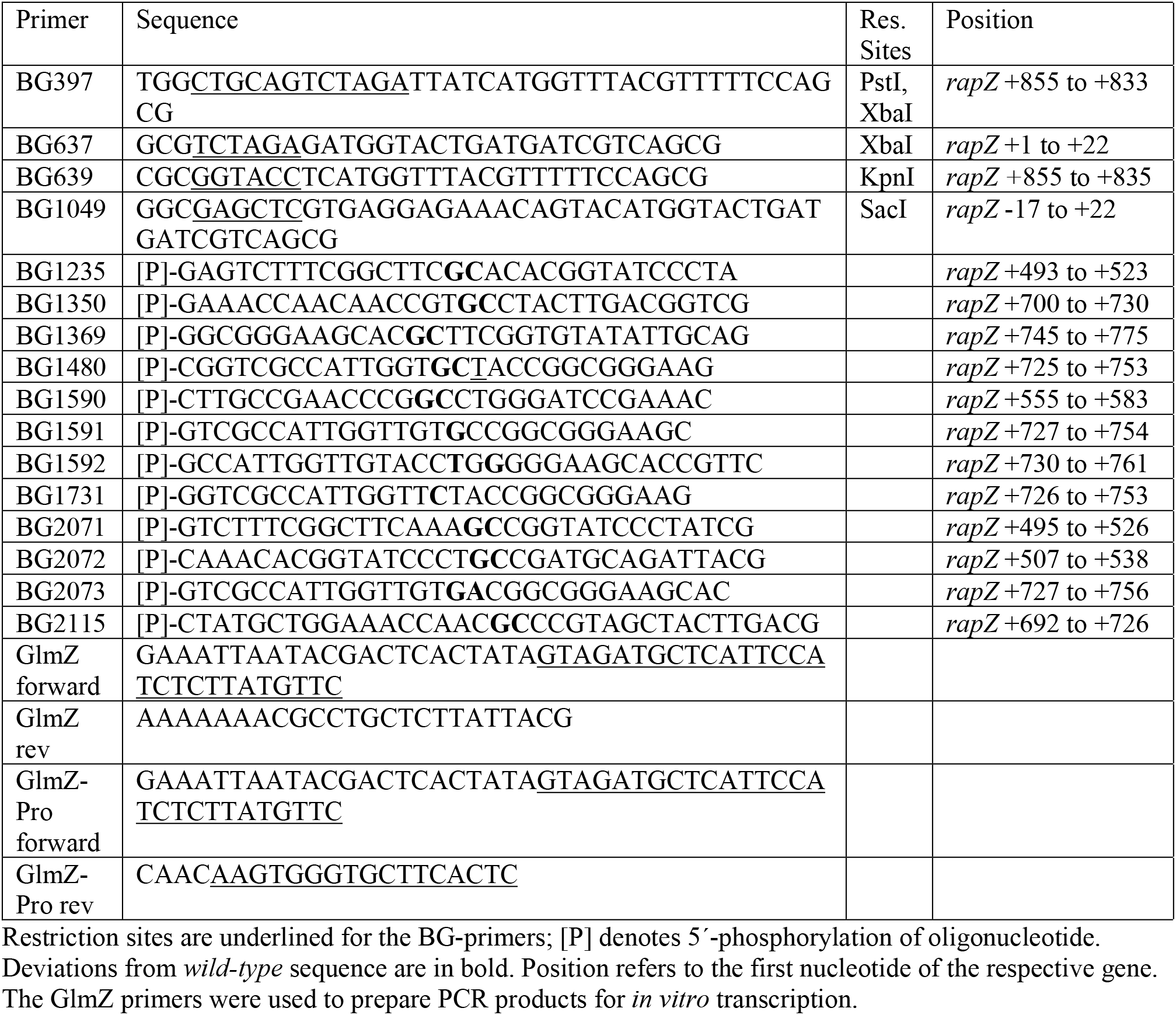
Oligonucleotides used in this study.

**Figure S1.**
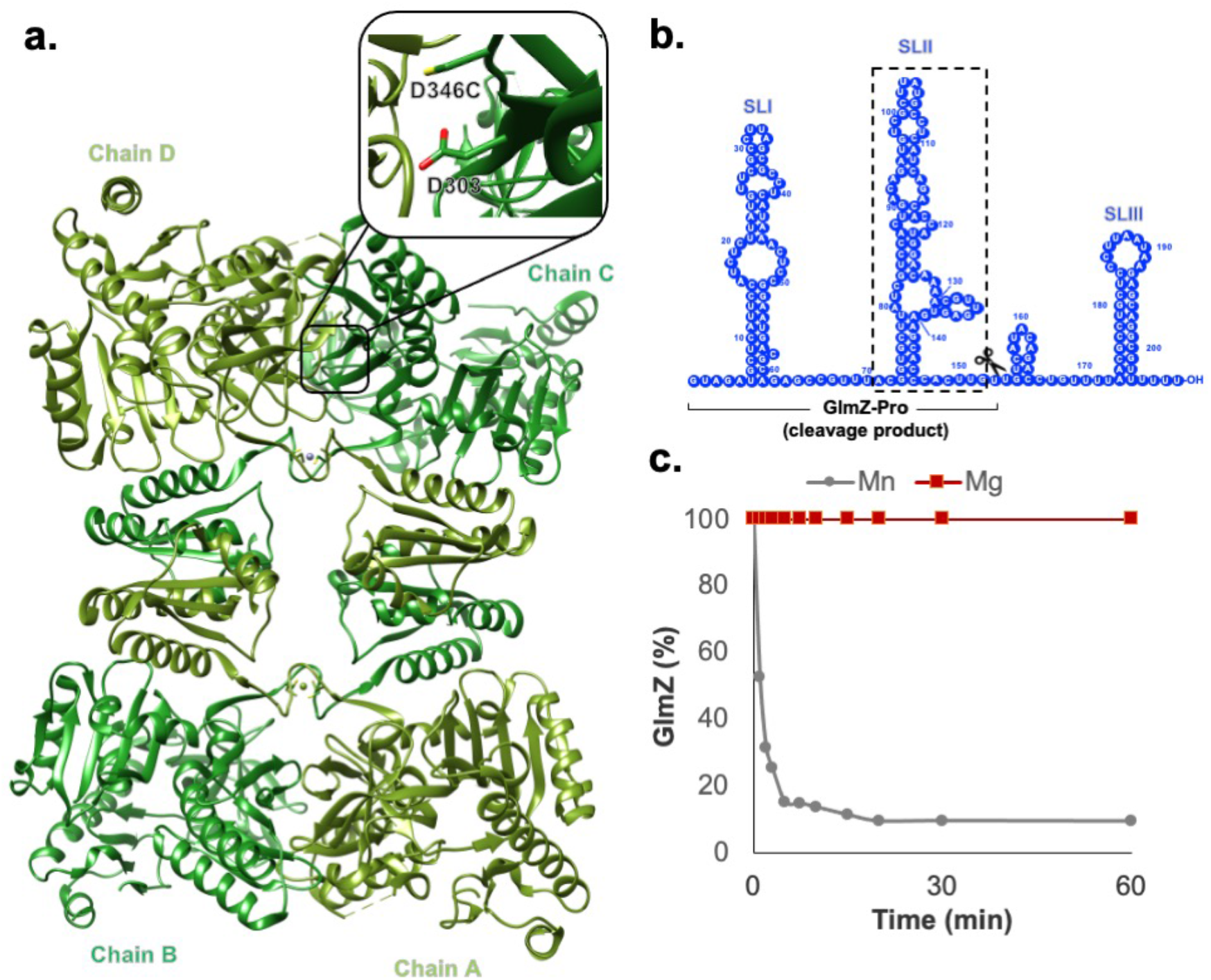
Metal-dependent nuclease activity of RNase E D346C. (a) Cartoon presentation of a crystal structure of RNase E-NTD in complex with RNA (PDB: 6G63). The RNA was omitted in the model for clarity. The depicted RNase E variant carries a substitution of residue Asp346 with Cys (D346C mutant). (b) RNA cleavage product detected in activity assays. (c) Activity assays confirm that the D346C mutant is inactivate with Mg^+2^ but highly active in the presence of Mn^+2^. Ribonuclease cleavage of GlmZ (625 nM) by RNase E-NTD (125 nM) in the presence of RapZ (250 nM) was carried out at 30° C in reaction buffer: 25 mM Tris-HCl pH 7.5, 50 mM NaCl, 50 mM KCl, 10 mM MgCl2 or MnCl2,1 mM DTT, 0.5 U/µL RNase OUT (Bandyra et al. 2018). Samples were quenched by adding proteinase K in 100 mM Tris-HCl pH 7.5, 150 mM NaCl, 12.5 mM EDTA, 1% SDS, followed by incubation at 50° C for 30 minutes. RNA samples were then mixed with loading dye (Thermo Fisher), heated at 95° C for 2 minutes and loaded onto 8% urea-PAGE gel. The gels were stained by SYBR^®^ Gold (ThermoFisher) and reaction products were visualized under UV transilluminator (GeneSnap, Syngene). To quantify, intensity of the reaction products was calculated using GeneTools (Syngene) against known amount of GlmZ.

**Figure S2.**
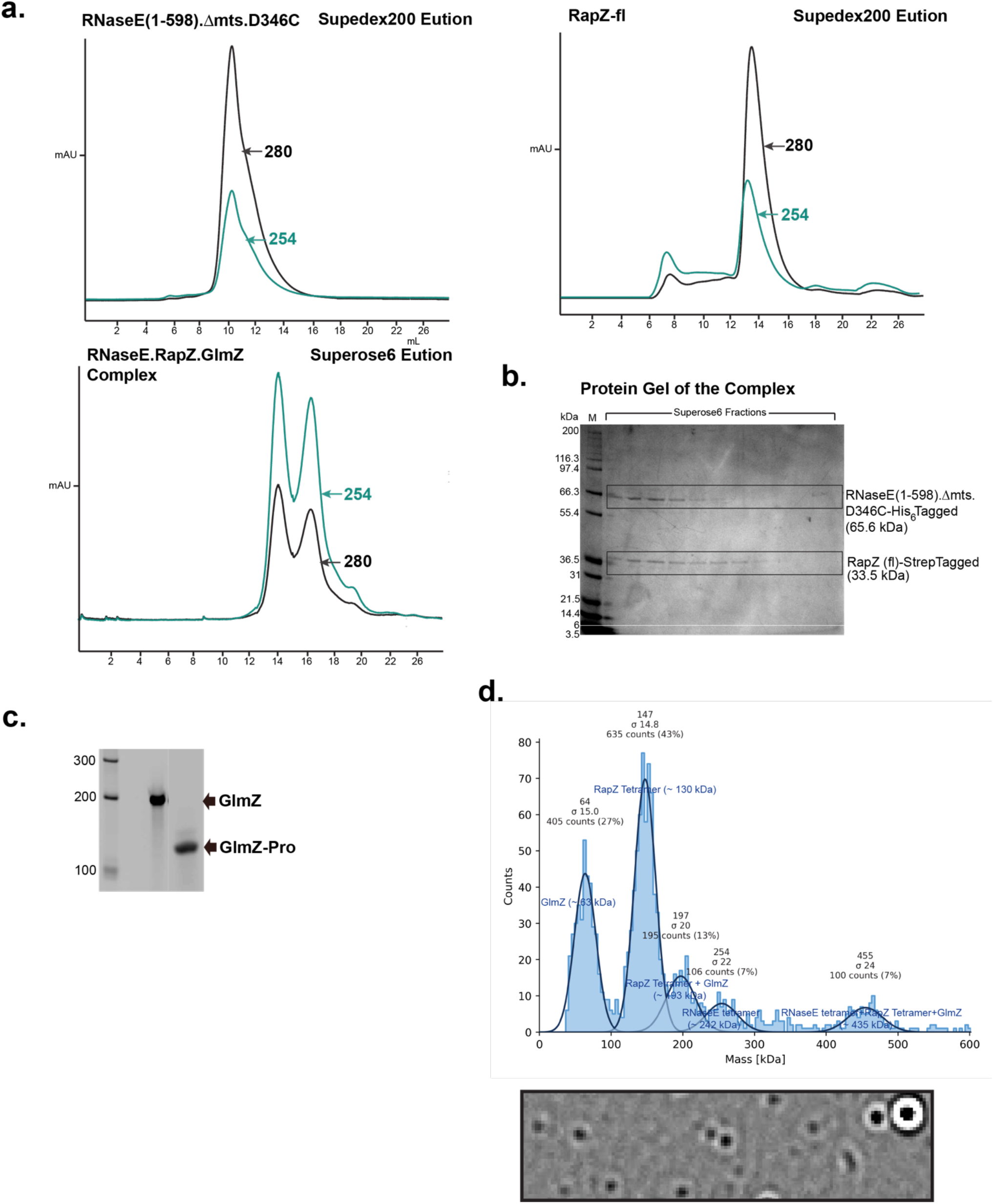
Gel filtration profile and mass photometry of the RNase E-NTD:GlmZ:RapZ complex. (a) Size-exclusion chromatograms showing a high 260/280 nm absorption ratio for the RNase E-NTD:RapZ:GlmZ complex compared to RNase E-NTD or RapZ; SDS-PAGE analysis confirming presence of RNase E and RapZ in the Superose 6 fractions. (b) Protein gel of the corresponding fractions from the SEC fractions. (c) Mass photometric analysis of the RNaseE-NTD:RapZ:GlmZ complex showing a heterogenous mixture. Mass Photometer Refeyn One (Refeyn Ltd., UK). The buffer used was 25 mM HEPES pH 7.5, 300 KCl, 1 mM MgCl2, 1mM DTT. (d) Denaturing RNA gel separating *in vitro* transcribed full-length GlmZ and processed GlmZ-pro for size comparison. The RNA size markers are annotated for length in nucleotides.

**Figure S3.**
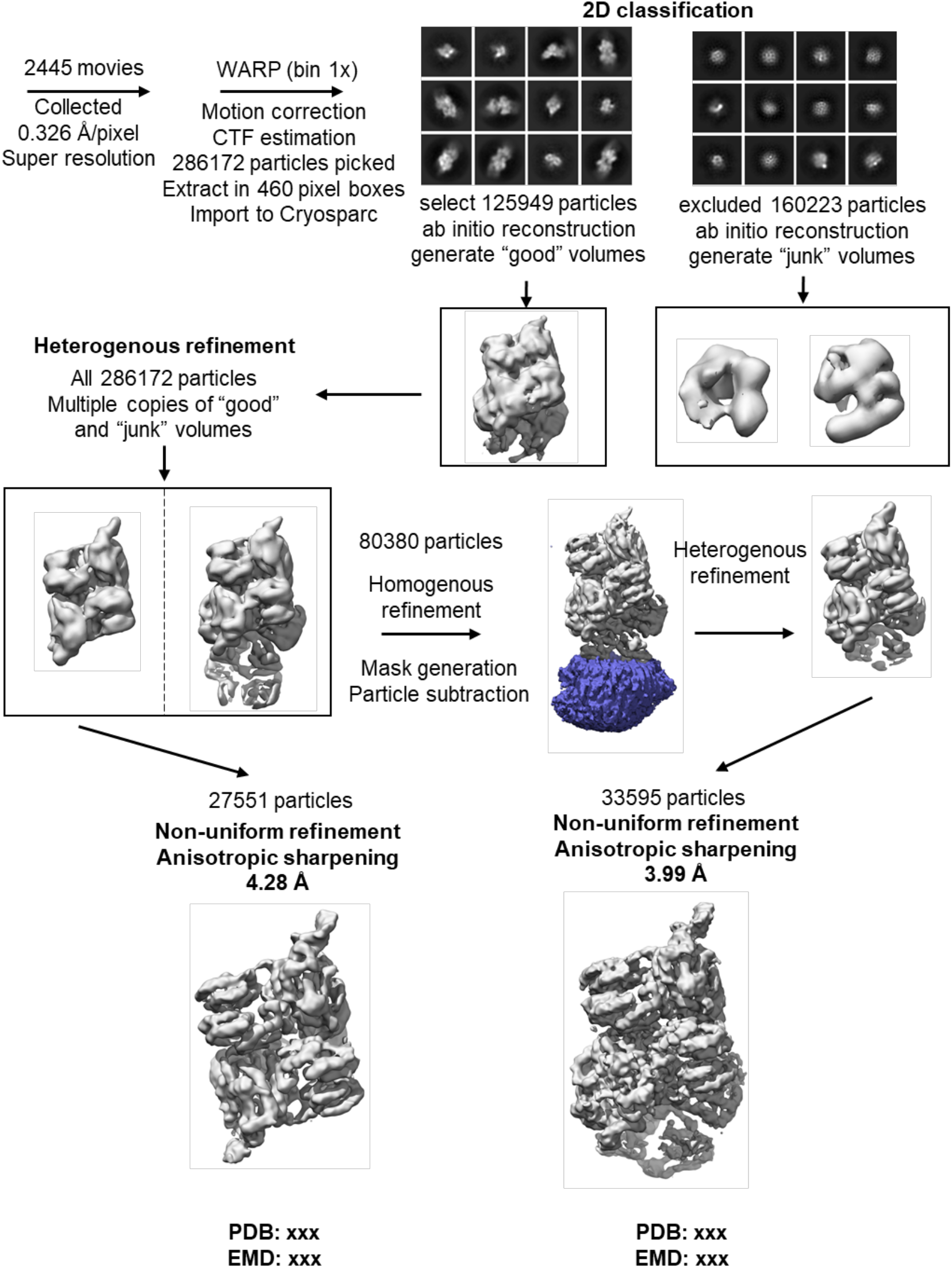
Workflow for data collection and processing. Initial grid preparation with quantifoil and ultrafoil grids showed aggregation or visible particles (not shown). Better results were obtained with graphene oxide covered grids.

**Figure S4.**
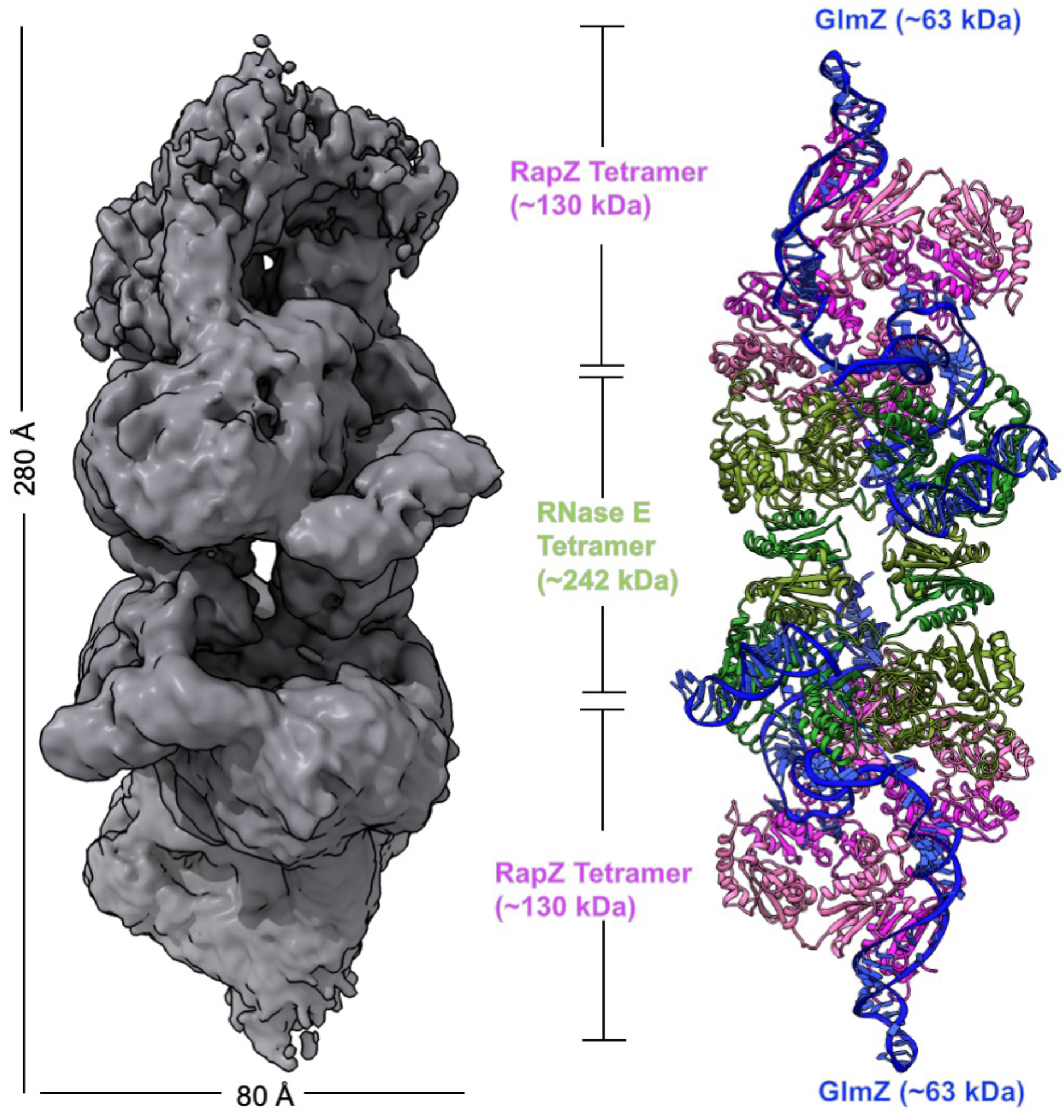
CryoEM model of the ternary RNase E-NTD:RapZ:GlmZ complex. The panel (left) shows the cryoEM map shown in grey. The model (right) of RNaseE-NTD:RapZ:GlmZ complex comprising one copy of RNase E-NTD tetramer (green), two copies of RapZ tetramer (pink), and two copies of GlmZ RNA (blue). Previously reported crystal structures of RapZ (PDB: 5O5O) and RNase E (PDB:6G63) and a model of GlmZ generated by ViennaRNA Package 2.0 were used to build the model of the ternary complex.

**Figure S5.**
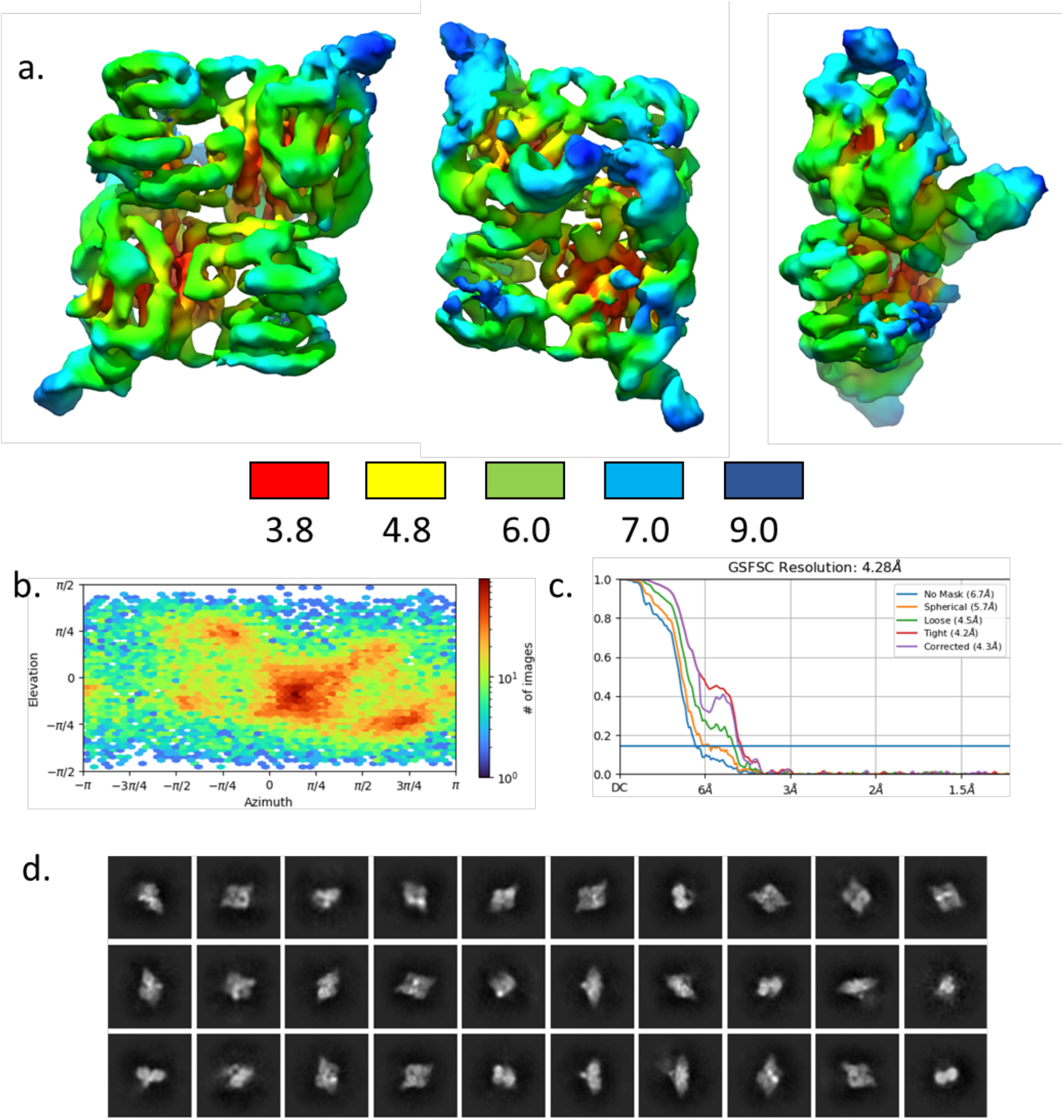
Summary of cryo-EM analysis of the RapZ:GlmZ binary complex. (a) Three views of the local resolution map of the binary complex calculated in cryoSPARC and coloured by resolution according to the key below. (b) Angular distribution calculated in cryoSPARC for particle projections contributing to the final map shown as a heat map. (c) Fourier schell correlation (FSC) resolution curves as calculated by cryoSPARC. (d) 2D class images from particles contributing to the final binary complex cryo-EM map.

**Figure S6.**
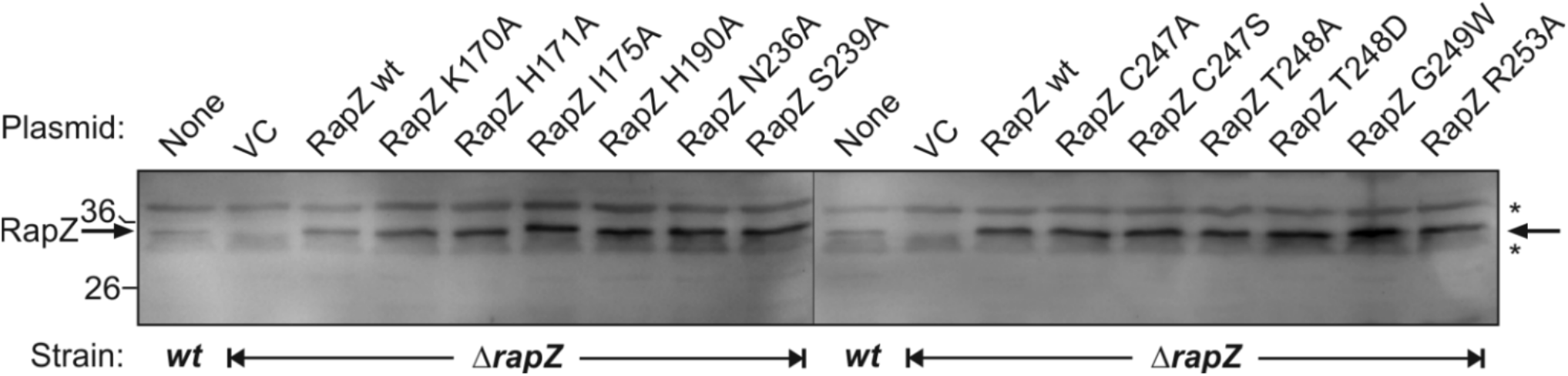
Western blot analysis verifying synthesis of plasmid-encoded RapZ variants. Total protein extracts of the arabinose-induced cultures tested in Figure 3b-c were separated by SDS-PAGE and gels were subsequently blotted. RapZ variants (indicated by arrow) were detected using a polyclonal antiserum against RapZ. Non-specifically detected proteins are indicated with asterisks.

**Figure S7.**
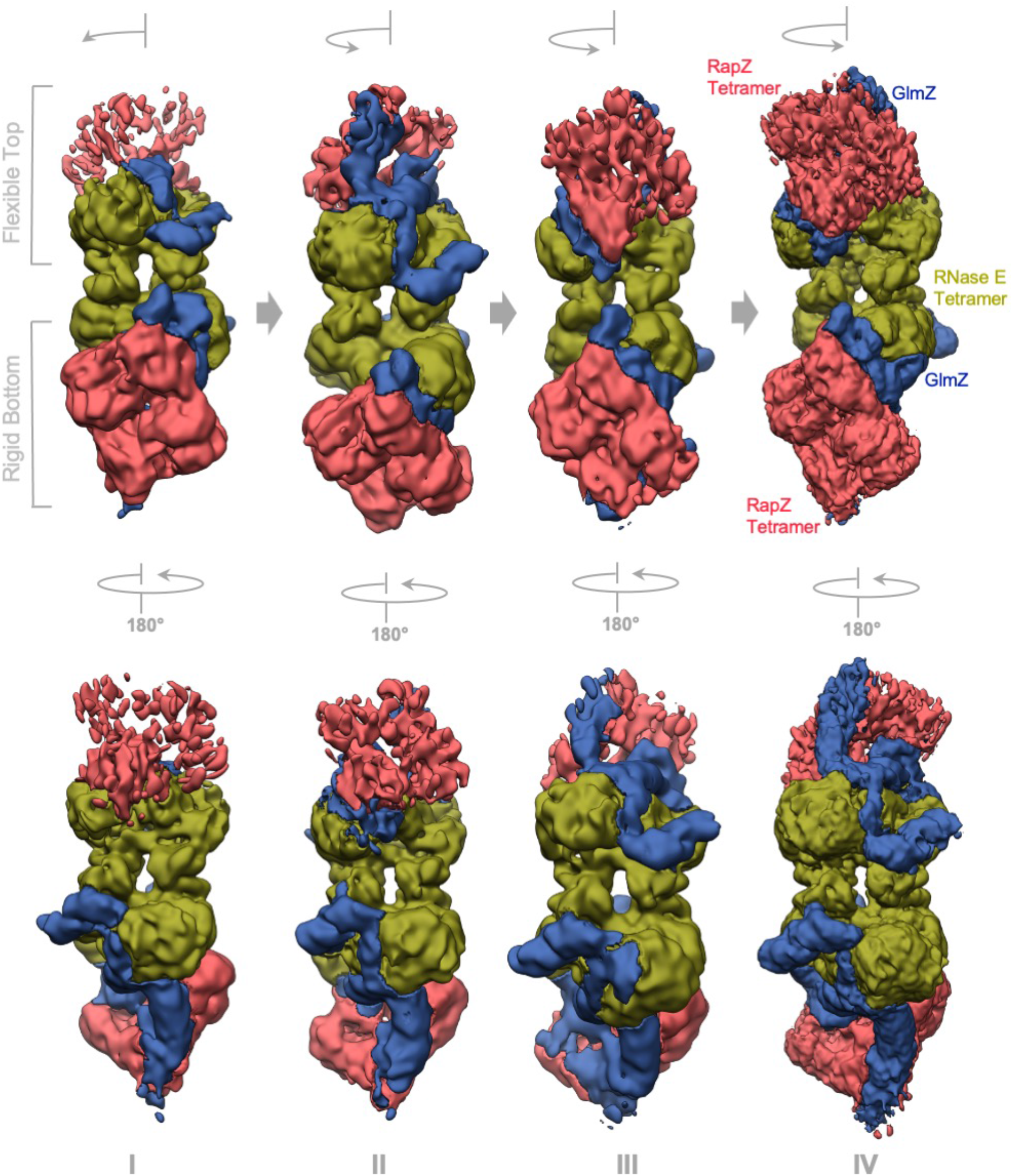
Heterogeneity of RNaseE-NTD:RapZ:GlmZ ternary complexes. Four assemblies (I-IV) of RNaseE-NTD:RapZ:GlmZ ternary complex revealing partial occupancy of RapZ:GlmZ binding to one of the RNase E-NTD dimers. Comparing classes II with III and IV, there is a *cis-/trans-*type binding of two RapZ tetramers with respect to each other on a RNaseE-NTD tetramer where one RapZ tetramer (bottom) is rigidly bound compared to the other which swivels around the apex of RNaseE-NTD tetramer.

**Figure S8.**
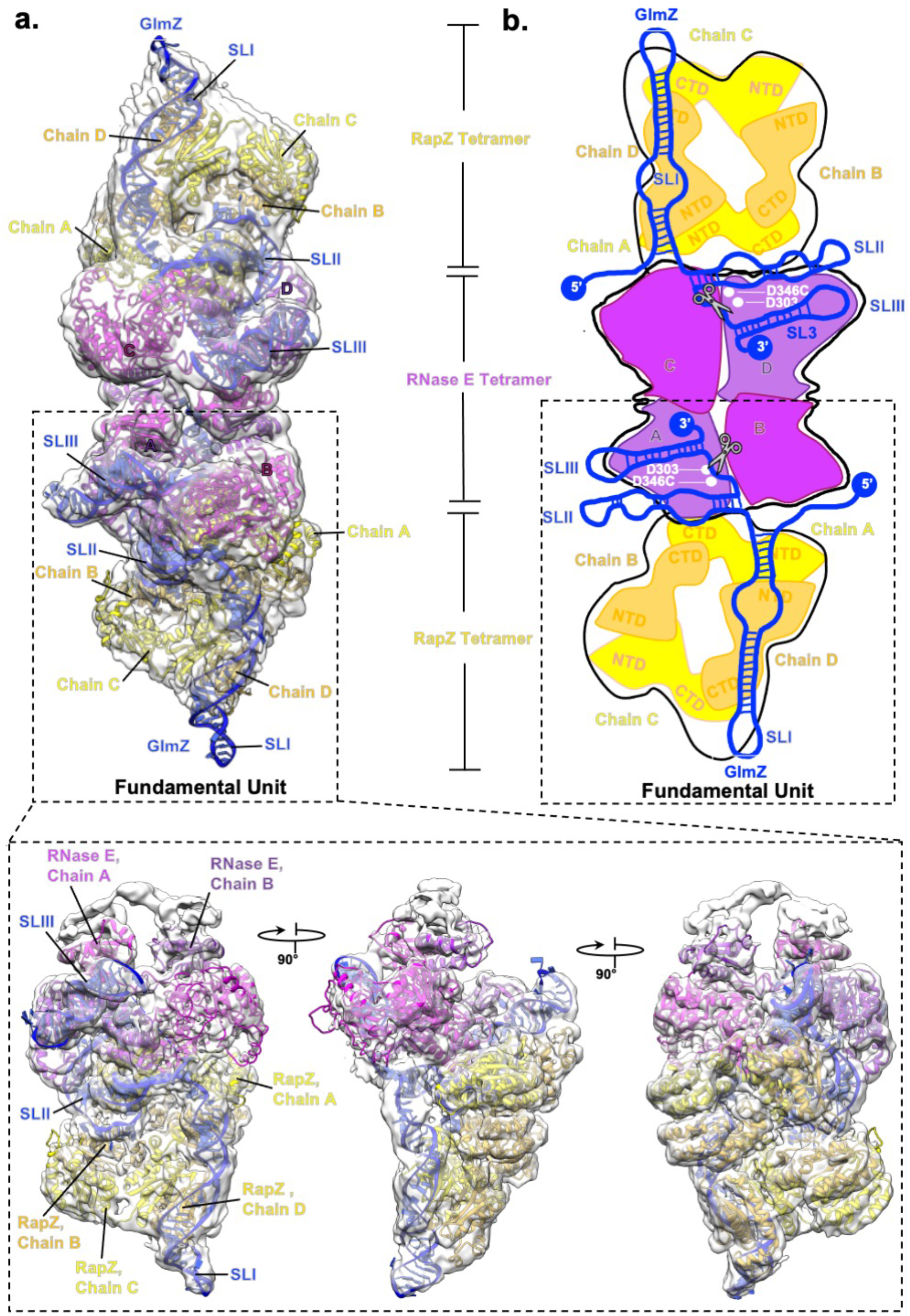
Fundamental unit of RNaseE-NTD:RapZ:GlmZ complex. (a) A rigid body fit of the model of the RNaseE-NTD:RapZ:GlmZ complex into the cryoEM map (white transparent) and (b) a cartoon representation of the model. A black box (dotted line) around the model and the cartoon indicates the fundamental unit of the complex; insets show orientation of the fundamental unit from three different angles.

**Figure S9.**
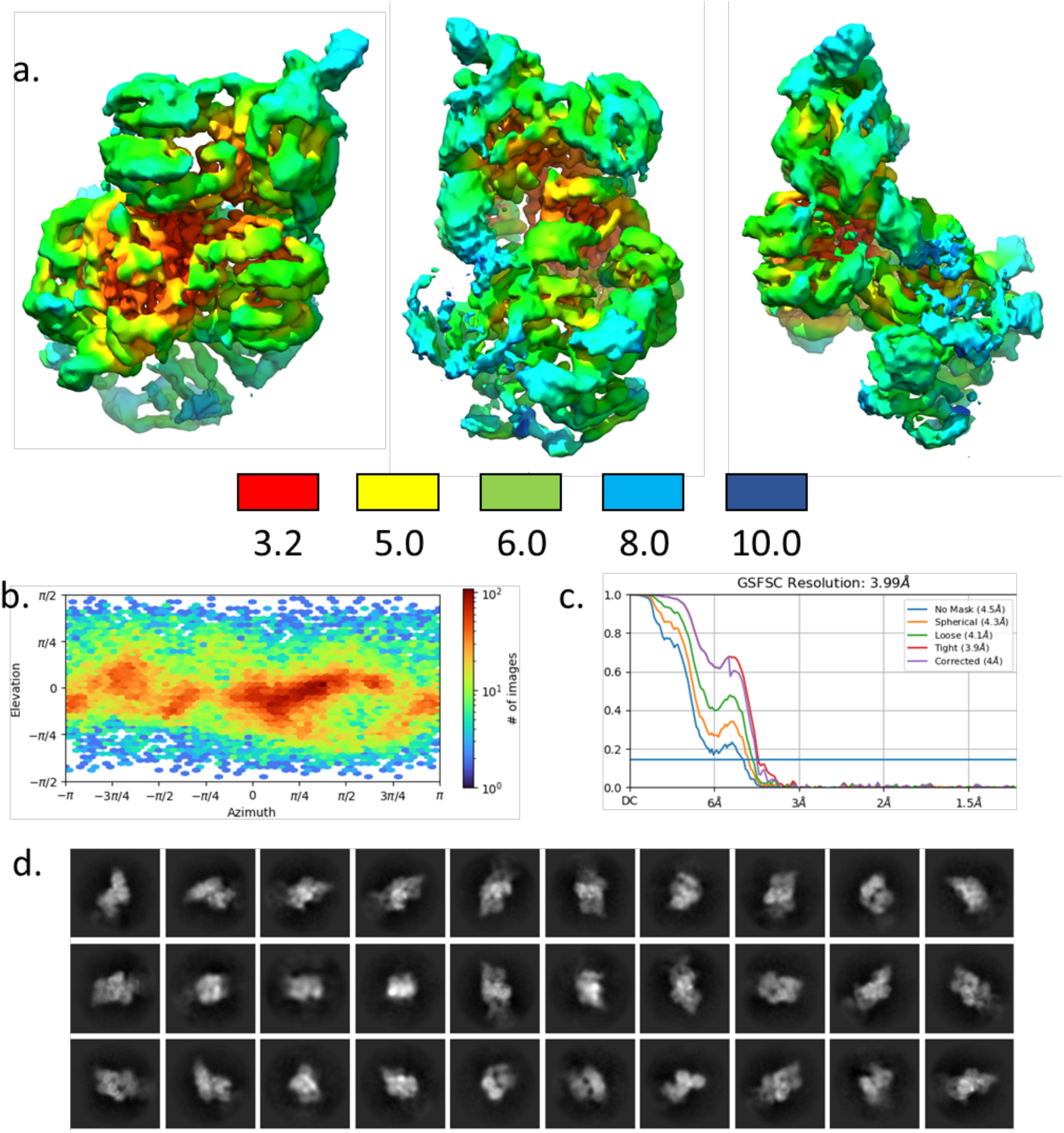
Summary of cryo-EM analysis of the RNase E-NTD:RapZ:GlmZ ternary complex. (a) Local resolution map of the ternary complex calculated in cryoSPARC and coloured by resolution according to the key below. (b) Angular distribution calculated in cryoSPARC for particle projections contributing to the final map shown as a heat map. (c) FSC resolution curves as calculated by cryoSPARC. **d)** 2D class images from particles contributing to the final ternary complex cryo-EM map.

**Figure S10.**
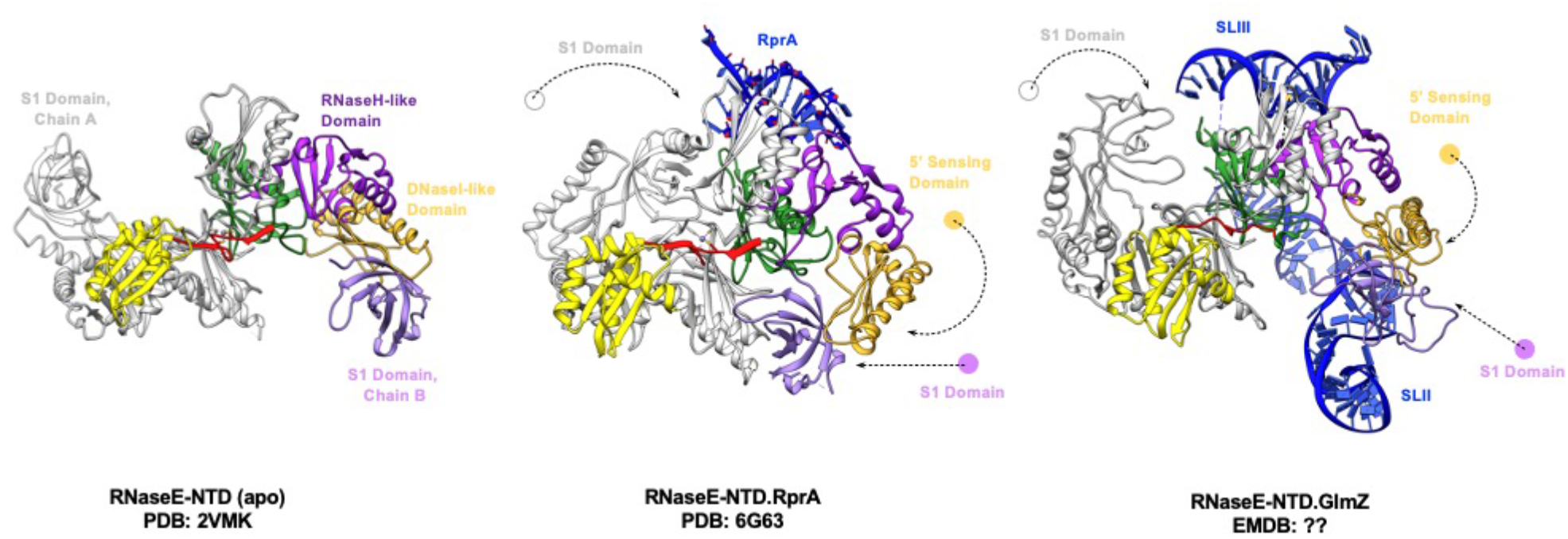
Conformational changes in the catalytic domain of RNase E. GlmZ-induced movement of the S1 (both catalytic and non-catalytic protomers), DNase I-like, and 5’-sensing domains; dotted arrows indicate the directionality of movements of the individual domain as compared with RNaseE-NTD apo-form. The red element is the Zn-mediated dimerization interface.

**Figure S11.**
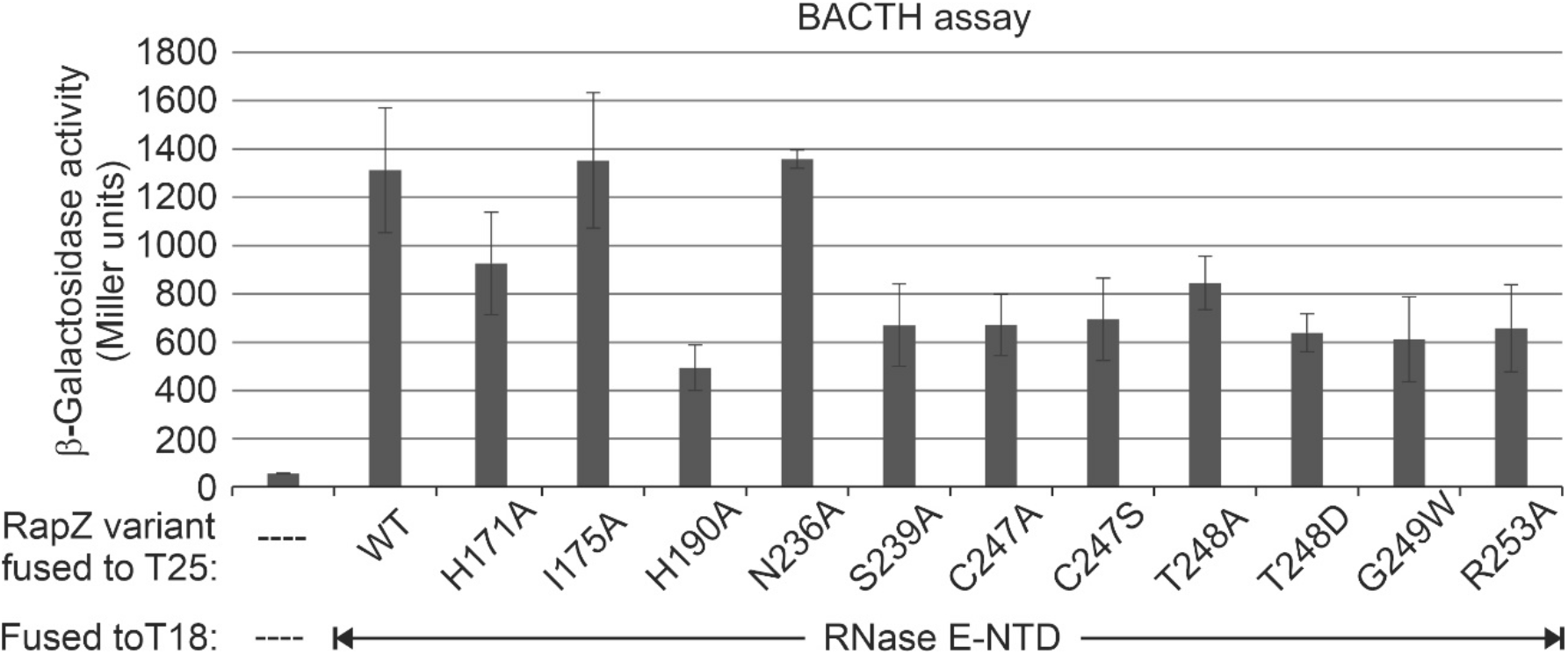
Bacterial adenylate cyclase-based two-hybrid (BACTH) assay to assess interaction performance of RapZ variants with the RNase E-NTD. BACTH measures efficiency of restoration of cAMP synthesis through interaction of proteins that are fused to split adenylate cyclase T18- and T25-fragments. Synthesis of endogenous β-galactosidase is measured as regulatory output. Reporter strain BTH101 lacks *cyaA* on the chromosome but carried two compatible plasmids encoding the T25- and T18-fragments either alone (pKT25/pUT18C vector control; column 1) or as fusion proteins with the indicated RapZ variant and the RNase E-NTD, respectively (cf. Table S2 for plasmids). Cells were grown to stationary phase in presence of 1 mM IPTG to allow for expression of the fusion genes from their plasmid-encoded *lac* promoters and for de-repression of the chromosomal *lac* operon. Average β-galactosidase activities including standard deviations are shown, which derive from at least three measurements and two independently transformed cell lines.

